# Adaptive metabolic strategies in consumer-resource models

**DOI:** 10.1101/385724

**Authors:** Leonardo Pacciani-Mori, Andrea Giometto, Samir Suweis, Amos Maritan

## Abstract

Bacteria are able to adapt to different environments by changing their “metabolic strategies”, i.e. the ways in which they uptake available resources from the environment. For example, in a celebrated experiment Jacques Monod showed that bacteria cultured in media containing two different sugars consume them sequentially, resulting in bi-phasic growth curves called “diauxic shifts”. From the theoretical point of view, microbial communities are commonly described using MacArthur’s consumer-resource model, which describes the population dynamics of species competing for a given set of resources. In this model, however, metabolic strategies are treated as constant parameters. Here, we introduce adaptive metabolic strategies in the framework of consumer-resource models, allowing the strategies to evolve to maximize each species’ relative fitness. By doing so, we are able to describe quantitatively, and without invoking any specific molecular mechanisms for the metabolism of the microbial species, growth curves of the baker’s yeast *Saccharomyces cerevisiae* measured in a controlled experimental set-up, with galactose as the primary carbon source. We also show that metabolic adaptation enables the community to self-organize, allowing species to coexist even in the presence of few resources, and to respond optimally to a time-dependent environment. A connection between the Competitive Exclusion Principle and the metabolic theory of ecology is also discussed.

## Introduction

Biodiversity is one of the most fascinating aspects of nature: from the microscopic to the continental scale, complex communities composed of tens to thousands of species compete for resources and yet co-exist. In particular, the survival of a species depends on the availability of resources in the environment, which is not static and can be altered by the presence of other species in the community. Furthermore, biodiversity is crucial for the functioning and maintenance of whole ecosystems, directly impacting their productivity, stability and many other properties [1]. The coexistence of several species in the same ecosystem can by analyzed and investigated experimentally using controlled microbial communities. Indeed, studies in the field of microbial ecology have shown that several species can coexist in the presence of few resources [2–5], and how this is possible is a long-standing open question [5–8].

Independent, and apparently unrelated, experimens of microbial batch growth have shown that microbes are capable of adapting to environments containing two or more resources [9–11]. Bacteria, in fact, are capable of adapting to different environments in various ways, by transforming and recycling nutrients and by varying the rates with which they uptake resources with time. Already in the early 1940s, Jacques Monod [9] observed that *Escherichia coli* and *Bacillus subtilis* grown in a culture medium containing two different sugars exhibit a bi-phasic growth curve, which he called “diauxie”. Instead of metabolizing these two nutrients simultaneously, bacteria consumed them sequentially using the most favorable one first (i.e., the one that conferred the highest growth rate) and once it had been depleted, following a lag phase, they resumed growth using the other sugar. Since then, diauxic growth has been the subject of thorough empirical study [12–15], via experiments that have generally involved the growth of one microbe on two resources, and the occurrence of diauxic shifts has been documented to occur widely across different microbial species [16–18]. It is generally thought [19] that the existence of diauxic shifts is “adaptive”, and the central idea of related modeling frameworks is that the regulatory processes behind diauxic shifts may be considered as the aftermath of an evolutionary optimization strategy [20]. Many models have been proposed to describe this phenomenon, but all are focused on the specific gene regulation and expression mechanisms of a given species [11, 21], and are generally tailored to describe the growth of such a species on a specific set of resources [19, 22, 23].

The pervasiveness and generality of these experimental results call for a theoretical framework capable of describing these phenomena in a unified way as emergent properties of complex systems of agents that interact with each other and with the environment, rather than through *ad hoc* tailored biological and/or molecular mechanisms. There is a growing effort from the Statistical Physics community to develop such a framework, with particular focus on the aspects of species coexistence and on the conditions leading to it [24–30]. The models devised in this direction typically build on MacArthur’s consumer-resource framework [31, 32], describing the competition of species for a common pool of resources, but neglecting microbial adaptation. In fact, despite the aforementioned evidence for adaptive metabolic strategies from studies of microbial metabolism, these models implicitly but systematically assume that the metabolic strategies, do not change with time, and assume that a species’ consumption rate of a given resource depends solely on the concentration of the latter, and not on the presence of other species, nor on the concentration of other nutrients. Goyal et al. [33] have recently developed a conceptual model of microbial communities that does not explicitly describe population dynamics, but where metabolic strategies can change.

Thus, even though it is clear that microbes can change the genetic expression level of different metabolic pathways in response to environmental cues, a connection between this phenomenon and consumer-resource ecological modeling, transcending the specificity of any particular microbial species and/or set of resources, is still missing.

In this work, we allow metabolic strategies to depend on time within a consumer-resource model, with dynamics that optimize the relative fitness of each species. This approach is capable of quantitatively reproducing experimentally-measured diauxic shifts. When considering multiple species and resources, our model suggests that adaptation plays a major role also in promoting species diversity, especially when few common resources are available. Furthermore, if the environmental conditions of the system are variable over time, or if some of the available resources degrade rapidly, our adaptive framework is capable of maintaining the coexistence of several species on few resources, while the “classic” MacArthur’s consumer-resource model would predict the extinction of most species in the system.

Therefore, our work proposes a unifying theoretical framework capable of reproducing both the existence of diauxic shifts and the coexistence of a large number of species competing for a limited number of resources in various realistic conditions.

## RESULTS

### The MacArthur’s consumer-resource model

In the classical formulation of MacArthur’s consumer-resource model, a community of *m* microbial species competes for *p* resources according to the following equations:

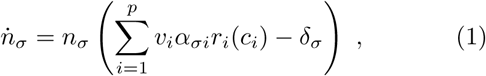

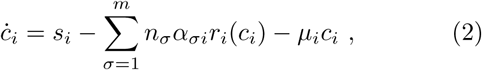

where *n*_*σ*_(*t*) describes the population density of species *σ, c*_*i*_(*t*) is the concentration of resource *i* and *δ*_*σ*_ is the death rate of species *σ*. The quantity *r*_*i*_(*c*_*i*_) is a function accounting for the fact that the dependence of a species’ growth rate on a given resource concentration saturates as *c*_*i*_ is increased. Without loss of generality, we assume that *r*_*i*_(*c*_*i*_) has the form of a Monod function [9], i.e. *r*_*i*_(*c*_*i*_) = *c*_*i*_*/*(*K*_*i*_ +*c*_*i*_) with *K*_*i*_ > 0, and so *r*_*i*_(*c*_*i*_) < 1 ∀*c*_*i*_ > 0. The quantities *α*_*σi*_ are the metabolic strategies, and each one of them can be interpreted as the maximum rate at which species *σ* uptakes resource *i*. The parameter *v*_*i*_ is often called “resource value” and is related to the resource-to-biomass conversion efficiency: the larger *v*_*i*_, the larger the population growth rate that is achieved for unit resource quantity, and thus the more “favorable” resource *i* is. The parameter *s*_*i*_ is a constant nutrient supply rate, and the sum in (2) represents the action of all consumers on resource *i*. Such an action depends of course on the metabolic strategies *α*_*σi*_. Finally, *µ*_*i*_ is the degradation rate of resource *i*.

### Introducing adaptive metabolic strategies

The introduction of dynamic metabolic strategies in the consumer-resource framework starts from the requirement that each metabolic strategy 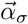 evolves in time to maximize the relative fitness of species *σ*, measured [34, 35] as the growth rate 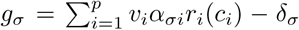. This can be achieved by requiring that metabolic strategies follow a simple gradient ascent equation:

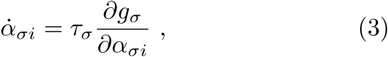

where 1*/τ*_*σ*_ is the characteristic timescale over which the metabolic strategy of species *σ* evolves.

Equation (3) is missing an important biological constraint, which is related to intrinsic limitations to any species’ resource uptake and metabolic rates: by necessity, microbes have limited amounts of energy that they can use to produce the metabolites necessary for resource uptake, so we must introduce such a constraint in (3). To do so, we require that each species has a maximum total resource uptake rate that it can achieve, i.e. 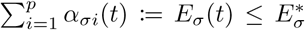. The choice of imposing a soft constraint in the form of an inequality is not arbitrary, as it is rooted in the experimental evidence that microbes cannot devote an unbounded amount of energy to metabolizing nutrients. Experiments [36] have shown, in fact, that introducing a constraint for metabolic fluxes in the form of an upper bound[37] allows one to improve the agreement between Flux Balance Analysis modeling and experimental data on *E. coli* growth on different substrates.

The constraint on the species’ maximum total resource uptake rates introduces a trade-off between the use of different resources. In the SI, we present a geometrical interpretation of the maximization problem given by (3), i.e. 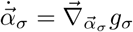 where 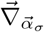 is the gradient with respect to the components of 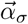. In particular, if we want 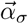 to evolve so that 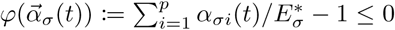, it is sufficient to remove from 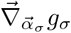 the component parallel to 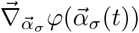 as soon as 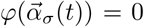. Furthermore, we prevent the metabolic strategies from becoming negative. Eventually, the final equation for the metabolic strategies’ dynamics is given by:

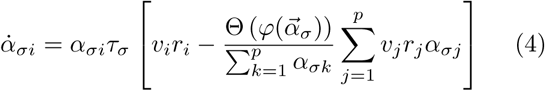

(see SI for the full derivation), where Θ is Heaviside’s step function (i.e. Θ(*x*) = 1 when *x ≥* 0 and Θ(*x*) = 0 otherwise). For the moment being, we assume that all the degradation rates *µ*_*i*_ are null, but we will later discuss a more general case.

### Diauxic shifts

If (4) is used alongside (1) and (2), the model is capable not only of reproducing qualitatively the growth dynamics of diauxic shifts, but to do so in quantitative agreement with experimental observations. To show this, we measured growth curves of the baker’s yeast, *S. cerevisiae*, grown in the presence of galactose as the primary carbon source. In these growth conditions, *S cerevisiae* partially respires and partially ferments the sugar. As a byproduct of fermentation, yeast cells release ethanol in the growth medium, which can then be respired by the cells once the concentration of galactose in the medium is reduced. To model the growth of *S. cerevisiae* in these conditions, we modified the equations to account for the fact that the second resource, ethanol, is produced by the yeast cells themselves, while the first one, galactose, is consumed (see SI for further details on the experiment and on the model equations). In Figure 1A, we show that our adaptive consumer-resource model can reproduce the experimental data with parameters that are compatible with values found in the literature (see Table S1). Figure 1B, on the other hand, shows that the “classic” MacArthur’s consumer-resource model with fixed metabolic strategies can surprisingly reproduce a diauxic-like behavior, but is incapable to describe quantitatively the experimental data as well as the adaptive model. Furthermore, one of the best fit parameters for the model with fixed metabolic strategies (*K*_*gal*_) is five orders of magnitude smaller than experimental values found in the literature (see Table S1), further suggesting that the model with fixed strategies cannot describe our experiment. The Akaike Information Criterion, used to compare the relative quality of the two models discounting the number of parameters, selects unambiguously the model with adaptive strategies as the best-fitting one (see SI for more information).

**FIG. 1:**
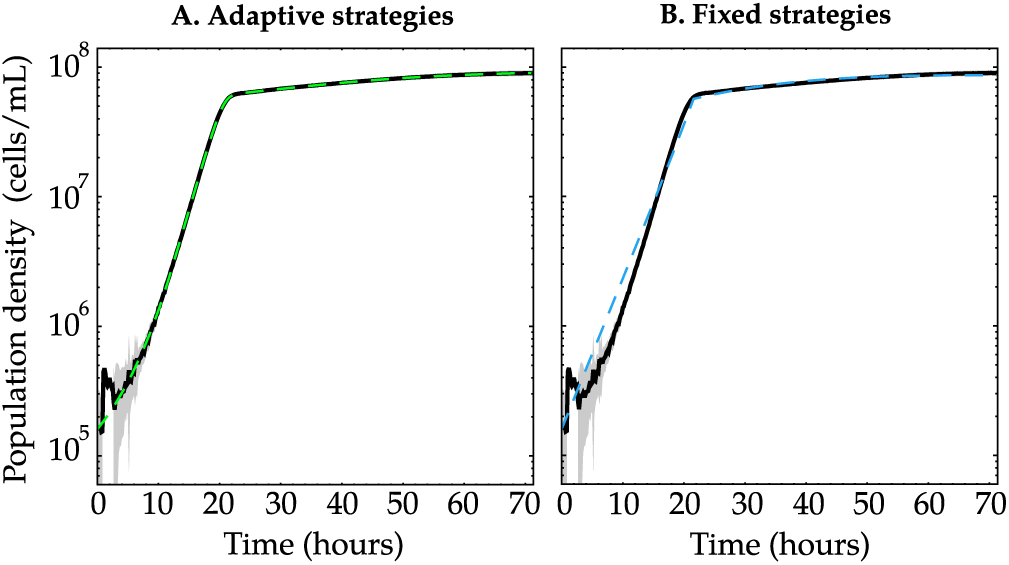
Comparison between the fits of MacArthur’s consumer-resource model (dashed lines) and experimental measures of the growth of *S cerevisiae* on galactose as the primary carbon source and ethanol as a byproduct of fermentation, in the case of adaptive (A) and fixed (B) metabolic strategies. Shown are the mean (black lines) and one standard deviation (gray bands) across *n* = 8 replicate populations. See SI for further details on the experiment, the model equations and the best-fit parameters.

### Species coexistence

The MacArthur’s consumer-resource model also makes predictions for the coexistence of *m* species on *p* shared resources and reproduces the so-called “Competitive Exclusion Principle” [38] (CEP), a theoretical argument that has sparked a lively debate in the ecological community [39–43]. According to the CEP, the maximum number of species that can stably coexist is equal to *p*. In nature, however, there are many situations in which the CEP appears to be violated: the most famous example of such violation is the “Paradox of the Plankton” [6], whereby a very high number of phytoplankton species is observed to coexist in the presence of a limited set of resources [44]. Many different mechanisms have been proposed to explain the violation of the CEP, ranging from non-equilibrium phenomena [6] to the existence of additional limiting factors like the presence of predators [45], cross-feeding relationships [5], toxin production [46], and complex or higher-order interactions [47, 48]; see [7, 8] for comprehensive reviews.

Considering now our model in the general case of *m* species and *p* resources, if the total maximum resource uptake rates 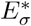 are completely uncorrelated to the death rates *δ*_*σ*_ (e.g. if 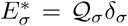, with 𝒬_*σ*_ drawn randomly from a given distribution with average ⟨𝒬⟩ and standard deviation Σ) we observe extinctions, i.e. in the infinite time limit, we cannot have more than *p* coexisting species (see SI). In Figure 2 we show how the time of first and seventh extinction changes as we vary the standard deviation Σ of the distribution from which we draw the 𝒬_*σ*_ in a system of *m* = 10 species and *p* = 3 resources. As shown, these extinction times increase sensibly as Σ is reduced, so the species present in the system can coexist for increasingly longer times, the more the 𝒬 _*σ*_ are peaked around their mean value. As we can see, the extinction times exhibit a power law-like behavior as a function of the coefficient of variation Σ*/*⟨𝒬⟩. We find that the times to extinction of the first *m* − *p* species scale approximately as (Σ*/*⟨𝒬⟩)^−1^. This observation suggests that coexistence for an *infinite* time interval could be possible if the ratio between the maximum resource uptake rate and the death rate of each species, which we call the Characteristic Timescale Ratio (CTR), does not depend on the species’ identities. In mathematical terms, if 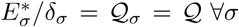, then *m* > *p* species can coexist. This requirement is compatible with the metabolic theory of ecology [49] (see SI for a detailed mathematical justification of this statement), according to which these two rates (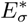 and *δ*_*σ*_) depend only on the characteristic mass of a species. It is indeed possible to show analytically that our model can violate the CEP if the CTR 𝒬 does not depend on the species’ identities (see SI). In this latter case, since a single time scale characterizes each species, we set *τ*_*σ*_ = *dδ*_*σ*_, where *d* regulates the speed of adaptation.

**FIG. 2:**
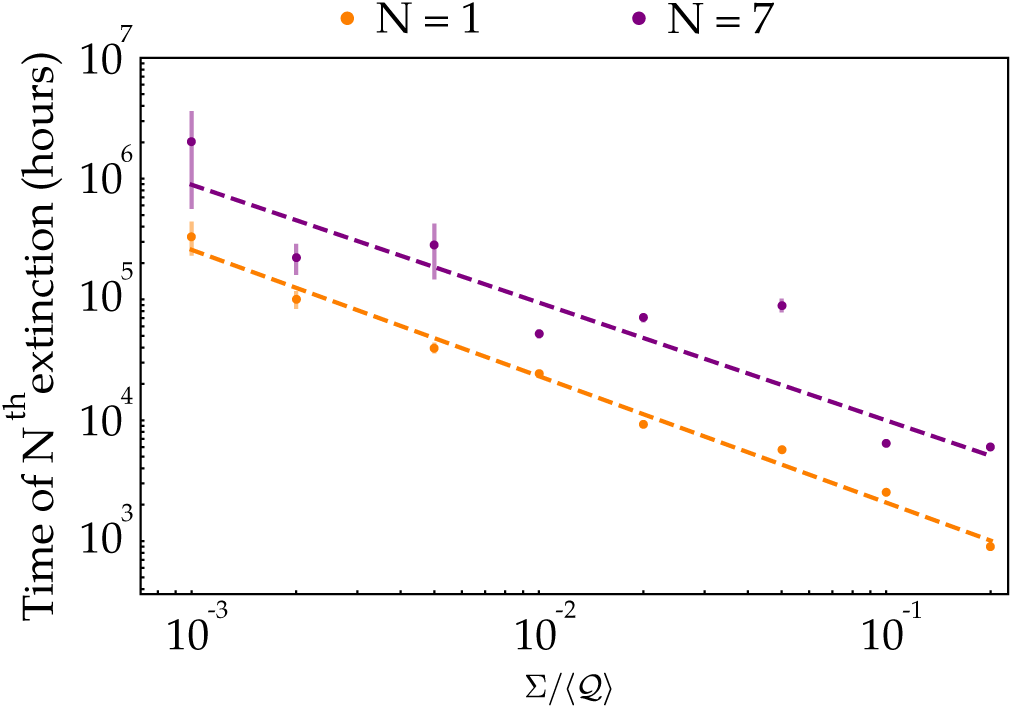
Time of first (orange) and seventh (purple) extinction in the consumer-resource model with adaptive metabolic strategies and 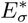 drawn independently of the *δ*_*σ*_. We used *m* = 10, *p* = 3 and 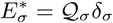 with 𝒬_*σ*_ drawn from a normal distribution with mean and ⟨𝒬⟩ standard deviation Σ; see SI for more details on the parameters used. The extinction times were computed as the instants at which the densities of the species fell below 1 cell/mL. Both axes are in logarithmic scale, the error bars represent one standard deviation across 200 iterations of the model and the dashed lines are the best power-law fits.

MacArthur’s consumer-resource model with fixed *α*_*σi*_ has been shown to violate the CEP only if 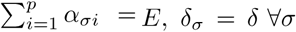 and 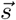 belongs to the convex hull of the metabolic strategies 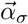 [25]. In general, any looser constraint (including 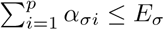 with arbitrary *E*_*σ*_) will lead to the extinction of at least *m* − *p* species, i.e. the system will obey the CEP; in this sense the system allows coexistence only when fine-tuned. However, if we now use (4) for the dynamics of *α*_*σi*_, it is possible to show analytically that the system gains additional degrees of freedom which make it possible to find steady states where an arbitrary number of species can coexist, even when the initial conditions are not favorable. More specifically, if we denote by 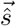 and 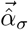 some appropriately rescaled versions of the nutrient supply rate 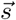 and the metabolic strategies 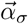 (see SI for more information), the system reaches a steady state where all species coexist even when 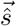 initially does not belong to the convex hull of 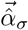. In Figure 3, we show the initial and final states of a temporal evolution of the model: even though 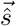 was initially outside the convex hull of 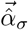, the metabolic strategies evolved to bring 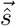 within the convex hull and thus allowed coexistence. Thus, the community modeled by Eqs. (1-4) is capable to *self-organize*. Notice that if we used fixed metabolic strategies in this case, almost all species would go extinct and the CEP would hold (see figure S2).

**FIG. 3:**
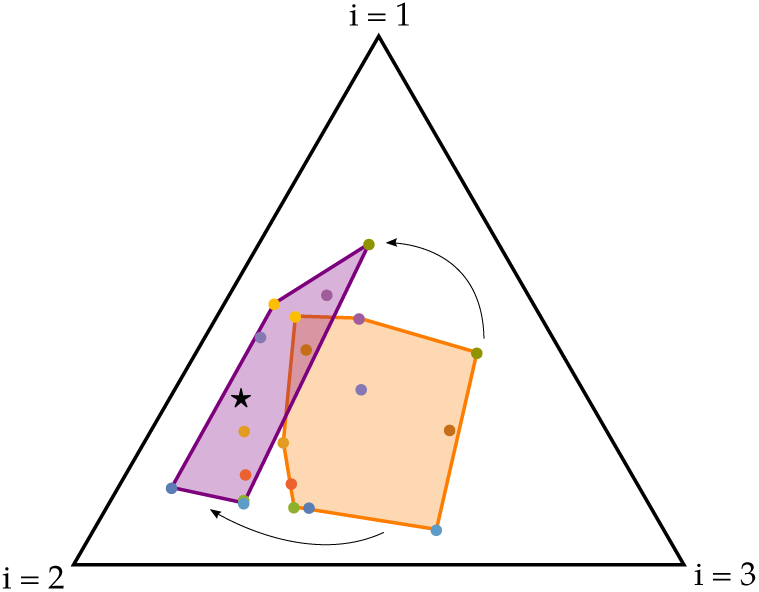
Comparison between the initial (orange) and final (purple) convex hull of the rescaled metabolic strategies (colored dots) when they are allowed to evolve according to (4). These results have been obtained for a system with *m* = 10 species and *p* = 3 resources using the graphical representation method introduced by Posfai et al. [25] and using a common value of the CTR 𝒬 for all species. Therefore, the rescaled metabolic strategies and nutrient supply rate vector (black star) all lie on a 2-dimensional simplex (i.e. the triangle in the figure), where each vertex corresponds to one of the resources; for details on the parameters used, and for the plots of the temporal evolution of the population densities and metabolic strategies, see Figure S3. In the final state, the 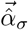 have incorporated 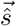 in their convex hull.

Therefore adaptation and the metabolic theory of ecology potentially provide one new fundamental mechanism for the coexistence of a large number of species on a limited number of resources.

#### Minimization of energy waste

An independent prediction of our model is that if one of the available resources, e.g. resource *j*, is too “energetically unfavorable”, then adaptation will bring all the *j*-th components of the metabolic strategies to zero, i.e. species will stop using that resource. The “unfavorableness” of resource *i* can be measured by 1*/v*_*i*_. When the metabolic strategies are not allowed to adapt, it is possible to prove that a nontrivial stationary state (i.e. one where the CEP is violated) is possible only if 1*/v*_*i*_ < 𝒬 ∀*i* ; this means that if even just one of the resources is unfavourable, i.e. 1*/v*_*j*_ > 𝒬 for one *j*, then there will be extinctions and in the end the CEP will hold (see SI and figure S4 for more details). However, when we allow the strategies to evolve following (4), the system reaches a non-trivial stationary state even if there is one (or possibly more) resource *j* for which 1*/v*_*j*_ > 𝒬. In this case, in fact, resource *j* becomes too unfavorable, and it is possible to show that the system “decouples” from that resource, i.e. the *j*-th component of *all* the metabolic strategies becomes null (see Figure S4). Something analogous happens also when degradation rates are present, i.e. *µ*_*i*_ > 0 in (2): in this case, at stationarity, the convex hull of the rescaled metabolic strategies will include the vector with components 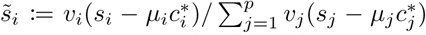 with 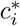 the stationary value of *c*_*i*_(*t*) (see SI), and if one of the *µ*_*i*_ is sufficiently large, this vector will lie on one of the sides of the (*p* − 1)-dimensional simplex where our system can be represented. In other words, we find that if the degradation rate *µ*_*j*_ of resource *j* becomes too large, then again all the *j*-th components of the metabolic strategies will become null (see Figures S5 and S7). On the other hand, if we introduce the resource degradation rates in MacArthur’s consumer-resource model with fixed metabolic strategies, extinctions will occur and the CEP will hold (see Figure S5) for any choice of *E*_*σ*_.

Therefore, species in our model minimize the energy they use to metabolize resources that are unfavorable or volatile, and they invest their energy budget on the more convenient ones.

#### Variable environmental conditions

Having adaptive metabolic strategies also allows the system to better respond to variable environmental conditions, i.e. when 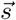 is a function of time 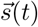. Let us consider a scenario where the nutrient supply rates change periodically; this can be implemented by shifting 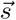 between two different values at regular time intervals: one inside the convex hull of the initial (rescaled) metabolic strategies and one outside of it. We found that when the metabolic strategies 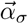 are allowed to evolve, the species’ populations oscillate between two values and manage to coexist, while when the metabolic strategies are fixed in time, some species go extinct due to the perturbations and the CEP is recovered, unless 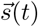 spends enough time inside the convex hull of the metabolic strategies – see Figure 4. Also in the case of environmental conditions that vary with time, we find that when we introduce non-null resource degradation rates that are sufficiently large, all the *i*-th components of the metabolic strategies vanish (see Figure S8). Therefore, adaptive metabolic strategies allow species in the community to efficiently deal with variable environmental conditions and a mix of (energetically) favorable and unfavourable resources, a characteristic feature of natural ecosystems.

**FIG. 4:**
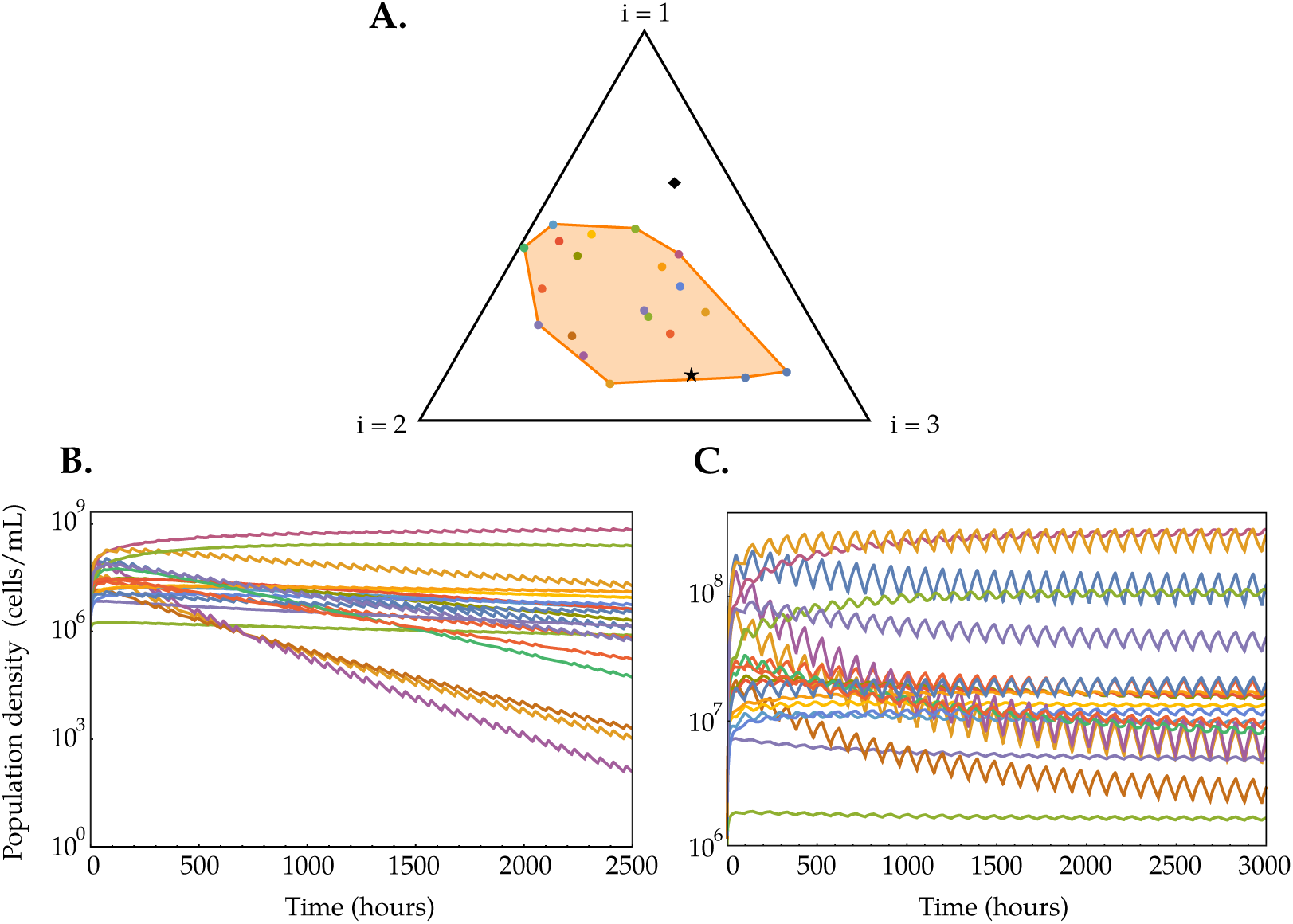
Comparison between the temporal dynamics of the population density of species *σ* = 20 in the consumer-resource models with fixed and adaptive metabolic strategies, when the resource supply rate vector 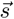 varies with time. Here, we simulated a system with *m* = 20 species, *p* = 3 resources, and with the nutrient supply rate vector switching at regular intervals between the two values shown (black star and diamond) in (A). Specifically, in (B) we made 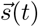 alternate periodically between 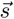 for *τ*_in_ = 12 h and 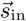 for *τ*_out_ = 48 h, with 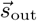 chosen within the convex hull of the initial rescaled metabolic strategies and 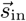 chosen outside it (see Figure S6 for more information on the parameters used). In (C) we have done the same, but with *τ*_in_ = *τ*_out_ = 48 h.

### Adaptation velocity

A physically relevant parameter characterizing the capacity of a species to adapt to a new environment is *d*, which regulates the adaptation velocity of the metabolic strategies (see (3) with *τ*_*σ*_ = *dδ*_*σ*_). Increasing the value of *d* leads to metabolic strategies that evolve more rapidly, and as a consequence species’ growth rates will be optimized for longer periods of time. Therefore, when *d* takes larger values, stationary population densities will be higher, while if *d* tends to zero we recover the case of fixed metabolic strategies and thus the CEP will determine the fate of the community. As shown in Figure 5, the distribution of stationary species’ populations can in-deed change sensibly with the adaptation velocity *d*, and if *d* is small enough then extinctions are observed and the CEP is recovered also in our model, so the number of available resource determines the limit of the system biodiversity.

**FIG. 5:**
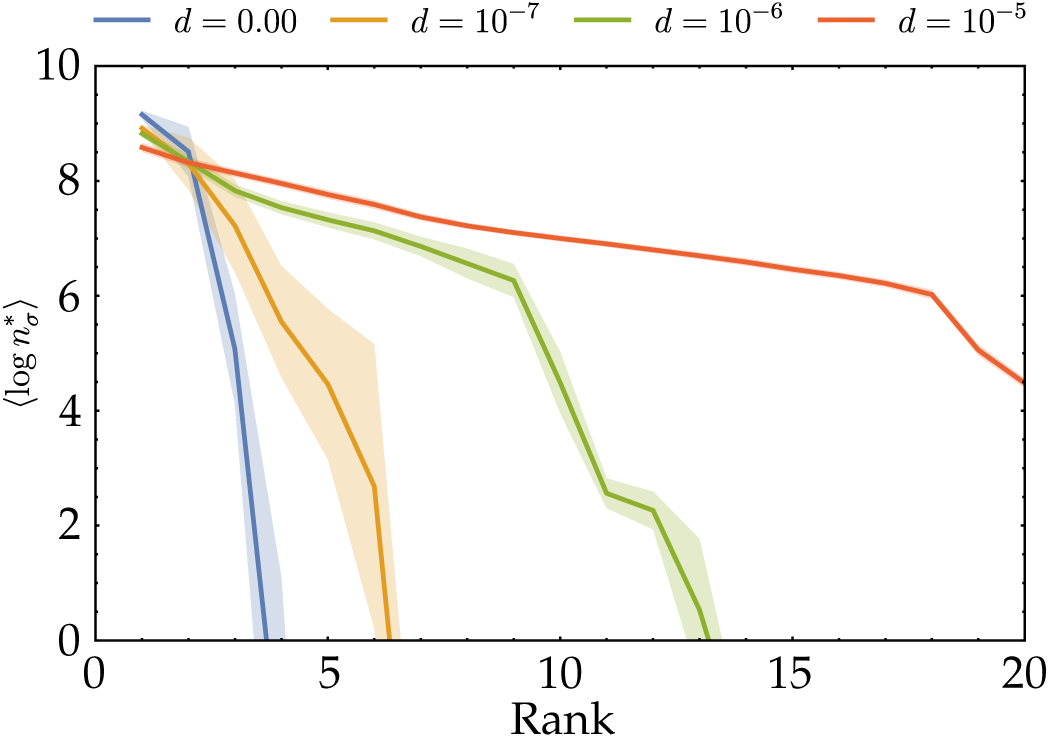
Rank distribution of the (decimal) logarithm of the stationary population densities 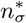 for different adaptation velocities (see Figure S9 for more information on the parameters used). The lines represent the average value over 100 iterations, while the opaque bands outline the standard error of the mean. As we can see, for *d* = 0 the rank distribution is very steep and only the first few species have a population density over 1 cell/mL (corresponding to log 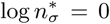), while as *d* increases the distribution becomes more even. If we set log 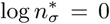 as our extinction threshold, we can see that with *d* = 10^−7^ approximately two thirds of the species in the system will go extinct, while with *d* = 10^−5^ all of them will survive.

On the other hand, if the system is subject to variable environmental conditions like the ones discussed in the previous paragraph (i.e. 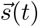 changes with time), as *d* increases the species’ are more able to promptly respond to perturbation and thus their populations will be less variable (see Figure S10).

## DISCUSSION

In this work we have introduced adaptive metabolic strategies, whose dynamics maximize each species’ growth rate, in MacArthur’s consumer-resource model. This new theoretical framework provides a unified description of phenomena, both at the individual and community species level, which were previously thought to be unrelated. Specifically, we have shown that consumer-resource models with adaptive metabolic strategies can quantitatively describe the growth of a single microbial species on multiple resources through a physical meaningful fit of the model parameters. Furthermore, we have found that the adaptive dynamics of metabolic strategies has a fundamental impact on species coexistence: the co-existence time interval of multiple species competing for few resources diverges as a characteristic timescale ratio (CTR) become less and less species-dependent. We suggest that this requirement is compatible with the metabolic theory of ecology [49], according to which the rates involved in the CTR depend only on the characteristic mass of the species. Without invoking the metabolic theory of ecology, each species would have its own CTR and extinctions would be unavoidable, leading to the CEP. Finally, we have shown that adaptation velocity is a crucial ingredient of our theoretical framework, such that CEP holds when adaptation is too slow with respect to the population dynamics.

In this work we thus propose a paradigmatic framework for consumer-resource models with adaptive metabolic strategies, by bridging the previously unrelated fields of microbial ecology and physiology and opening new theoretical paths ahead.

Although we have focused only on competitive interactions, these are clearly not the only kind of interactions found in natural communities. Recent studies have shown that phenomena such as cross-feeding and syntrophy are ubiquitous in the microbial world [50], and have crucial roles in shaping the structure and function of microbial communities [5, 29, 51]. A version of MacArthur’s consumer-resource model with cross-feeding dynamics and fixed metabolic strategies has been recently proposed [5]. Future work will be dedicated to generalizing our framework for systems with cross feeding. Recently, it has been observed experimentally that natural microbial communities are often composed of metabolically distinct and interdependent groups of species, each specialized in a particular function [2, 3, 52–54]; this property has also been tackled theoretically [55]. Another natural development of the framework presented here would be to investigate if this “modular” organization is an emergent property of the system and how such modularity influences the dynamics of species’ population abundances and the properties of the community.

## Supporting information

Animation of figures 3 and S3

Animation of figure S4

Animation of figure S5

Animation of figure S6a and S6b

Animation of figure S6a and S6d

## ACKNOWLEDGMENTS

We are grateful to J. Grilli for useful discussions. A. M. and L. P.-M. acknowledge the Cariparo Foundation, S. S. acknowledges the University of Padua for SID2017 and STARS2018 grants. A. G. was supported by research fellowships from the Swiss National Science Foundation, Projects P2ELP2 168498 and P400PB 180823.

