## Supplementary Information for "Adaptive metabolic strategies in consumer-resource models"

### Supplementary Information for
### "Adaptive metabolic strategies in consumer-resource models"

Leonardo Pacciani-Mori, Samir Suweis, and Amos Maritan*
*Department of Physics and Astronomy "Galileo Galilei"*
*University of Padua*
*Via Francesco Marzolo 8, 35131 Padua (Italy)*

Andrea Giometto
*Department of Physics*
*Department of Molecular and Cellular Biology*
*Harvard University*
*52 Oxford St, Cambridge, MA 02138*

#### CONTENTS

I. Some analytical results on the model 1
A. Results 1
B. Resource decoupling 3
C. Dimensional analysis 4

II. Adaptive metabolic strategies 4
A. Constrained equation for the metabolic strategies 5
B. Comparison between the model and experimental measures of diauxic growth curves 6
1. Methods 6
2. Results 7

III. Coexistence of multiple species 7
A. Extinction times 7
B. Metabolic theory of ecology 7
C. Unfavorable resources 8
D. Variable environmental conditions 8
E. Slow adaptation can still lead to competitive exclusion 9

References 9

#### I. SOME ANALYTICAL RESULTS ON THE MODEL

##### A. Results

We first consider our consumer-resource model with fixed metabolic strategies (we write $r_i$ instead of $r_i(c_i)$ from now on for the sake of simplicity):

$$\dot{n}_\sigma = n_\sigma \left( \sum_{i=1}^{p} v_i \alpha_{\sigma i} r_i - \delta_\sigma \right) , \tag{S.1a}$$

$$\dot{c}_i = s_i - \sum_{\sigma=1}^{m} n_\sigma \alpha_{\sigma i} r_i - \mu_i c_i . \tag{S.1b}$$

*

For the moment, we set $\mu_i = 0$. At stationarity, we have $\dot{n}_\sigma = 0\ \forall\sigma$ and $\dot{c}_i = 0\ \forall i$, so:

$$\sum_{i=1}^{p} \alpha_{\sigma i} v_i r_i^* = \delta_\sigma \ , \tag{S.2a}$$

$$r_i^* = \frac{s_i}{\sum_{\sigma=1}^{m} n_\sigma^* \alpha_{\sigma i}} \ , \tag{S.2b}$$

where we have used the symbol "*" to denote that a quantity is computed at stationarity. (S.2a) is a system of $m$ equations in $p$ unknowns ($r_i^*$) that, if $m > p$, won't admit nontrivial solutions unless extremely particular conditions hold. For example, we could require that the metabolic strategies are subject to some kind of constraint, e.g. $\sum_{i=1}^{p} \alpha_{\sigma i} = E_\sigma$, and it is easy to see that a possible solution of (S.2a) is:

$$r_i^* = \frac{1}{v_i} \cdot \frac{\delta_\sigma}{E_\sigma} \ , \tag{S.3}$$

which is impossible since the left hand side of (S.3) depends only on the index $i$ while the right hand side depends also on $\sigma$. In this case, therefore, it is impossible to find conditions under which all species coexist in the stationary state.

As stated in the Main Text and shown in Fig. 2, however, the model suggest that coexistence of all the species is possible if, in agreement with the metabolic theory of ecology, we set $E_\sigma^* = \mathcal{Q}\delta_\sigma$ (see section III B for a detailed discussion of this point). If we introduce this assumption, then (S.2a) becomes solvable even when $m > p$, and in particular we obtain:

$$r_i^* = \frac{1}{v_i}\mathcal{Q}^{-1} \ . \tag{S.4}$$

Therefore, since $r_i^* < 1$, we must have $1/v_i < \mathcal{Q}\ \forall i$, i.e. $v_i > \mathcal{Q}^{-1}\ \forall i$. This means that the benefit derived from using resources, as measured by $v_i$, must be advantageous enough: if this is not the case, the energy available for growth after the metabolization of a resource would be insufficient for growth, which from a biological point of view would be completely inefficient.

If we now introduce

$$\hat{n} \coloneqq \sum_{\sigma=1}^{m} n_\sigma \delta_\sigma \qquad\qquad n \coloneqq \sum_{\sigma=1}^{m} n_\sigma \ , \tag{S.5}$$

then, summing both sides of (S.1a) over $\sigma$ we obtain:

$$\dot{n} = \sum_{\sigma=1}^{m}\sum_{i=1}^{p} n_\sigma v_i \alpha_{\sigma i} r_i - \hat{n} \ , \tag{S.6}$$

which at stationarity gives $\hat{n}^* = \sum_{\sigma=1}^{m} n_\sigma^* \delta_\sigma = \sum_{i=1}^{p} v_i s_i$, because from (S.1b) we have:

$$s_i = \sum_{\sigma=1}^{m} n_\sigma^* \alpha_{\sigma i} r_i^* \ . \tag{S.7}$$

If we now use (S.4) and define:

$$x_\sigma^* \coloneqq \frac{n_\sigma^* \delta_\sigma}{\sum_{\rho=1}^{m} n_\rho^* \delta_\rho} \qquad\qquad \hat{s}_i \coloneqq \frac{v_i s_i}{\sum_{j=1}^{p} v_j s_j} \qquad\qquad \hat{\alpha}_{\sigma i} \coloneqq \frac{\alpha_{\sigma i}}{\mathcal{Q}\delta_\sigma} \ , \tag{S.8}$$

we have $\sum_{\sigma=1}^{m} x_\sigma^* = 1$, $\sum_{i=1}^{p} \hat{s}_i = 1$ and $\sum_{i=1}^{p} \hat{\alpha}_{\sigma i} = 1\ \forall\sigma$ (so both $\vec{\hat{s}}$ and $\vec{\hat{\alpha}}_\sigma$ belong to a $(p-1)$-dimensional simplex), and (S.7) can be rewritten as:

$$\hat{s}_i = \sum_{\sigma=1}^{m} x_\sigma^* \hat{\alpha}_{\sigma i} \ . \tag{S.9}$$

This is a system of $p$ equations for $m$ unknowns ($x_\sigma^*$), so if $m > p$ it will admit *infinite* solutions; however, to have species coexistence at stationarity and violate the Competitive Exclusion Principle we need (S.9) to admit a *positive*

solution $x^*_\sigma > 0\ \forall\sigma$. We also have $\sum_{\sigma=1}^m x^*_\sigma = 1$, so coexistence will be possible only if $\vec{\hat{s}}$ is a convex combination of the rescaled metabolic strategies $\vec{\hat{\alpha}}_\sigma$; in other words, the Competitive Exclusion Principle can be violated for an arbitrary number of species if $\vec{\hat{s}}$ belongs to the *convex hull* of the rescaled metabolic strategies $\vec{\hat{\alpha}}_\sigma$.
It is important to notice that consumer-resource models like this exhibit an anomalous case: if $p = 1$ and $m > 1$, in fact, (S.9) becomes

$$\hat{s} = \sum_{\sigma=1}^m x^*_\sigma \hat{\alpha}_\sigma \ , \tag{S.10}$$

where $\hat{s} = 1$, $\hat{\alpha}_\sigma > 0\ \forall\sigma$, so there are *infinite* possible solutions. In this model, therefore, when only one resource is present *all* competing species will always coexist, while when $p > 1$ resources are supplied to the system competitive exclusion can happen if $\vec{\hat{s}}$ does not belong to the *convex hull* of the rescaled metabolic strategies $\vec{\hat{\alpha}}_\sigma$.

Now, if we allow the metabolic strategies to evolve (see section II for details on how to derive the equations for $\alpha_{\sigma i}$) then (S.9) will be substituted by:

$$\hat{s}_i = \sum_{\sigma=1}^m x^*_\sigma \hat{\alpha}^*_{\sigma i} \ . \tag{S.11}$$

This is a system of $p$ equations for $(m+1)p$ unknowns ($x^*_\sigma$ and $\hat{\alpha}^*_{\sigma i}$) and thus *always* admits *infinite* solutions. In other words even if the initial conditions do not satisfy (S.9), the populations and metabolic strategies will evolve so that in the end the equation holds, and thus all species will coexist since their rescaled metabolic strategies will have "incorporated" $\vec{\hat{s}}$ in their convex hull.

##### B. Resource decoupling

Here, we study what happens when 1) there is one $j$ for which $1/v_j > \mathcal{Q}$ and when 2) the degradation rates are non-null.

1) If $1/v_j > \mathcal{Q}$ for one $j$, (S.4) does not have a feasible solution for $i = j$ and we can rewrite (S.2a) as:

$$\delta_\sigma = v_j\alpha^*_{\sigma j}r^*_j + \sum_{i\neq j} v_i\alpha^*_{\sigma i}r^*_i = v_j\alpha^*_{\sigma j}r^*_j + \sum_{i\neq j}\alpha^*_{\sigma i}\mathcal{Q}^{-1} = v_j\alpha^*_{\sigma j}r^*_j + \mathcal{Q}^{-1}\left(\sum_{i=1}^p \alpha^*_{\sigma i} - \alpha^*_{\sigma j}\right)\ \forall\sigma \qquad , \tag{S.12}$$

and since $\sum_{i=1}^p \alpha^*_{\sigma i} = \mathcal{Q}\delta_\sigma$ this reduces to:

$$\alpha^*_{\sigma j}(v_j r^*_j - \mathcal{Q}^{-1}) = 0\ \forall\sigma\ . \tag{S.13}$$

However, since (S.4) does not have a solution for $j$, the only way to solve (S.13) is to have $\alpha^*_{\sigma j} = 0\ \forall\sigma$: all the $j$-th components of the metabolic strategies are null at stationarity, i.e. all the species will stop using resource $j$. This means that resource $j$ "decouples" from the system, i.e. all species will act as if the $j$-th resource does not even exist; as a consequence, at stationarity the metabolic strategies will have changed so that their rescaled versions $\vec{\hat{\alpha}}_\sigma$ have incorporated the vector $\vec{\hat{s}}$ with $\hat{s}_j = 0$. Notice also that since $\alpha^*_{\sigma j} = 0\ \forall\sigma$, from (S.1b) we will have that after a transient $\dot{c}_j = s_j$, i.e. the concentration of resource $j$ grows linearly in time.
Notice that if the metabolic strategies $\alpha_{\sigma i}$ are fixed and do not change over time and $1/v_j > \mathcal{Q}$ for one $j$, since (S.4) does not have a feasible expression ($r^*_j > 1$) we cannot find solutions of (S.2a), and so ultimately we cannot find steady states where all species coexist. This means that in MacArthur's consumer-resource model with fixed metabolic strategies having even just one unfavourable resource will bring several species to extinction until the CEP holds.
Figure S.4 shows the results of a numerical simulation of the model in this case.

2) If we let $\mu_i \neq 0$, at stationarity (S.1b) leads to:

$$s_i - \mu_i c^*_i = \sum_{\sigma=1}^m n^*_\sigma \alpha^*_{\sigma i} r^*_i \ , \tag{S.14}$$

instead of (S.7). If we define:

$$\tilde{s}_i := \frac{v_i(s_i - \mu_i c_i^*)}{\sum_{j=1}^{p} v_j(s_j - \mu_j c_j^*)} \ , \tag{S.15}$$

in the end we will have:

$$\tilde{s}_i = \sum_{\sigma=1}^{m} x_\sigma^* \hat{\alpha}_{\sigma i}^* \ , \tag{S.16}$$

i.e. the rescaled metabolic strategies at stationarity will have changed so that $\vec{\tilde{s}}$ lies inside their convex hull. Notice that from (S.15) the rescaled nutrient supply rates depend also on the stationary values $c_i^*$ of the resources' concentrations.

Figure S.5 shows the results of a numerical simulation of the model in this case.

##### C. Dimensional analysis

Here, we perform a dimensional analysis of our model, i.e. (S.1a) and (S.1b). Since the $n_\sigma$s are population densities, we have $[n_\sigma]$ = population/volume = cells/mL, and given that the $c_i$s are resource concentrations, we have $[c_i]$ = resource/volume = g of resource/mL; furthermore, $[\delta_\sigma]$ = 1/time = 1/h. The resource availabilities $r_i$ are dimensionless, i.e. $[r_i] = 1$, and from (S.1b) we have:

$$\frac{[c_i]}{\text{time}} = [n_\sigma][\alpha_{\sigma i}] \qquad \Rightarrow \qquad [\alpha_{\sigma i}] = \frac{\text{g of resource}}{\text{cells} \cdot \text{h}} \ , \tag{S.17}$$

so the metabolic strategies represent indeed the amount of resource uptaken per cell per hour. From (S.1a) we have:

$$\frac{1}{\text{time}} = [v_i][\alpha_{\sigma i}] \qquad \Rightarrow \qquad [v_i] = \frac{\text{cells}}{\text{g of resource}} \ . \tag{S.18}$$

Furthermore, since $\sum_i \alpha_{\sigma i} \le \mathcal{Q}\delta_\sigma$ we have $[\alpha_{\sigma i}] = [\mathcal{Q}][\delta_\sigma]$ and so:

$$[\mathcal{Q}] = \frac{[\alpha_{\sigma i}]}{[\delta_\sigma]} = \frac{\text{g of resource}}{\text{cells}} \ . \tag{S.19}$$

Finally, $[d] = 1$ and $[s_i]$ = g of resource/mL · h.

#### II. ADAPTIVE METABOLIC STRATEGIES

As stated in the Main Text, we want each $\vec{\alpha}_\sigma$ to evolve so that the growth rate of species $\sigma$,

$$g_\sigma = \sum_{i=1}^{p} v_i \alpha_{\sigma i} r_i - \delta_\sigma \ , \tag{S.20}$$

is maximized. This can be achieved by simply requiring that the metabolic strategies evolve according to the "gradient ascent" dynamics:

$$\dot{\alpha}_{\sigma i} = d\delta_\sigma \frac{\partial g_\sigma}{\partial \alpha_{\sigma i}} = v_i r_i d\delta_\sigma \ . \tag{S.21}$$

This equation alone, however, does not prevent $\alpha_{\sigma i}$ from growing indefinitely; as stated in the Main Text, we must introduce a constraint on the metabolic strategies because microbes have limited amounts of energy they can use for resource uptake.

##### A. Constrained equation for the metabolic strategies

Let us therefore see how (S.21) changes if we introduce some constraints; we use now a completely general formalism, and then apply the results to the cases we are interested in. Let $Q(\vec{x})$ be the quantity that we want to maximize with the temporal evolution of a variable $\vec{x}(t) \in \mathbb{R}^q$, and suppose that during this evolution the variable $\vec{x}(t)$ is subject to a constraint $\varphi(\vec{x}) = 0$. A temporal evolution of the form

$$\dot{\vec{x}} = \vec{\nabla} Q(\vec{x}) \tag{S.22}$$

(which is the equivalent of (S.21) in our framework) implies, by scalarly multiplying both sides by $\dot{\vec{x}}$, that

$$\dot{\vec{x}}^2 = \vec{\nabla} Q(\vec{x}) \cdot \dot{\vec{x}} = \dot{Q} \ , \tag{S.23}$$

so that indeed $\dot{Q} \geq 0$. To keep $\varphi(\vec{x})$ constant, we simply eliminate the component of $\vec{\nabla} Q$ parallel to $\vec{\nabla}\varphi$, i.e.

$$\dot{\vec{x}} = \vec{\nabla} Q(\vec{x}) - \frac{\vec{\nabla}\varphi(\vec{x})}{|\vec{\nabla}\varphi(\vec{x})|} \left( \frac{\vec{\nabla}\varphi(\vec{x})}{|\vec{\nabla}\varphi(\vec{x})|} \cdot \vec{\nabla} Q(\vec{x}) \right) \ . \tag{S.24}$$

Figure S.1 contains an illustrative representation of this procedure.
Let us now show that this equation satisfies our requirements. First of all, $\varphi$ is constant along the trajectories:

$$\dot{\varphi} = \vec{\nabla}\varphi \cdot \dot{\vec{x}} = \vec{\nabla}\varphi \cdot \vec{\nabla} Q - \frac{|\vec{\nabla}\varphi|^2}{|\vec{\nabla}\varphi|^2} \vec{\nabla}\varphi \cdot \vec{\nabla} Q = 0 \ , \tag{S.25}$$

and $Q$ increases with time:

$$\dot{Q} = \vec{\nabla} Q \cdot \dot{\vec{x}} = |\vec{\nabla} Q|^2 - \left( \frac{\vec{\nabla}\varphi}{|\vec{\nabla}\varphi|} \cdot \vec{\nabla} Q \right)^2 \geq 0 \ , \tag{S.26}$$

which follows from Schwartz's inequality: $(\vec{a} \cdot \vec{b})^2 \leq |\vec{a}|^2 |\vec{b}|^2$, and so $(\vec{\nabla}\varphi \cdot \vec{\nabla} Q / |\vec{\nabla}\varphi|)^2$ can never be greater than $|\vec{\nabla} Q|^2$. Of course, if the constraint has the "softer" form $\varphi(\vec{x}) \leq 0$, then the final equation for $\vec{x}(t)$ becomes:

$$\dot{\vec{x}} = \vec{\nabla} Q(\vec{x}) - \Theta(\varphi(\vec{x})) \frac{\vec{\nabla}\varphi(\vec{x})}{|\vec{\nabla}\varphi(\vec{x})|} \left( \frac{\vec{\nabla}\varphi(\vec{x})}{|\vec{\nabla}\varphi(\vec{x})|} \cdot \vec{\nabla} Q(\vec{x}) \right) \ , \tag{S.27}$$

where $\Theta$ is the Heaviside step function, i.e. $\Theta(x) = 0$ if $x \leq 0$ and $\Theta(x) = 1$ if $x > 0$. If we now rewrite (S.24) with the substitutions $\vec{x} \to \vec{\alpha}_\sigma$ (we can in fact suppose in general that the constraint depends on all metabolic strategies) and $Q \to d\delta_\sigma g_\sigma$, we obtain:

$$\dot{\alpha}_{\sigma i} = d\delta_\sigma \frac{\partial g_\sigma}{\partial \alpha_{\sigma i}} - \frac{\partial \varphi(\vec{\alpha}_\sigma)/\partial \alpha_{\sigma i}}{\sum_{\tau=1}^m \sum_{k=1}^p \left( \partial \varphi(\vec{\alpha}_\sigma)/\partial \alpha_{\tau k} \right)^2} \sum_{\rho=1}^m \sum_{j=1}^p \frac{\partial \varphi(\vec{\alpha}_\sigma)}{\partial \alpha_{\rho j}} d\delta_\rho \frac{\partial g_\sigma}{\partial \alpha_{\rho j}} \ . \tag{S.28}$$

This equation, however, does not guarantee in general that metabolic strategies remain non-negative (the second term on the right hand side of (S.28) could be larger than the first and lead $\alpha_{\sigma i}$ to negative values); we must therefore modify it so that $\alpha_{\sigma i}(t) \geq 0 \ \forall t$. The simplest way to do so is to introduce some auxiliary variables $\eta_{\sigma i}$ of which the metabolic strategies are some non-negative functions, i.e.

$$\alpha_{\sigma i} := \mathcal{F}(\eta_{\sigma i}) \qquad \text{with} \qquad \mathcal{F}(x) \geq 0 \ \forall x \ , \tag{S.29}$$

and then use (S.28) as an equation for $\eta_{\sigma i}$, i.e. we write:

$$\dot{\eta}_{\sigma i} = d\delta_\sigma \frac{\partial g_\sigma}{\partial \eta_{\sigma i}} - \frac{\partial \varphi(\vec{\eta}_\sigma)/\partial \eta_{\sigma i}}{\sum_{\tau=1}^m \sum_{k=1}^p \left( \partial \varphi(\vec{\eta}_\sigma)/\partial \eta_{\tau k} \right)^2} \sum_{\rho=1}^m \sum_{j=1}^p \frac{\partial \varphi(\vec{\eta}_\sigma)}{\partial \eta_{\rho j}} d\delta_\rho \frac{\partial g_\sigma}{\partial \eta_{\rho j}} \ . \tag{S.30}$$

If we now change variables and write this equation in terms of $\alpha_{\sigma i}$, we obtain:

$$\dot{\alpha}_{\sigma i} = \mathcal{F}'(\eta_{\sigma i})^2 \left[ d\delta_\sigma \frac{\partial g_\sigma}{\partial \alpha_{\sigma i}} - \frac{\partial \varphi(\vec{\alpha}_\sigma)/\partial \alpha_{\sigma i}}{\sum_{\tau=1}^m \sum_{k=1}^p \left( \mathcal{F}'(\eta_{\tau k}) \partial \varphi(\vec{\alpha}_\sigma)/\partial \alpha_{\tau k} \right)^2} \sum_{\rho=1}^m \sum_{j=1}^p \frac{\partial \varphi(\vec{\alpha}_\sigma)}{\partial \alpha_{\rho j}} d\delta_\rho \frac{\partial g_\sigma}{\partial \alpha_{\rho j}} \mathcal{F}'(\eta_{\rho j}) \right] \ . \tag{S.31}$$

We are now free to choose any form for $\mathcal{F}$. We have made two simple choices corresponding to $\mathcal{F}(x) = x^2/4$ (the 1/4 factor has been chosen so that the final equation for $\alpha_{\sigma i}$ doesn't have superfluous numerical factors) and $\mathcal{F}(x) = e^x$. The results for these two case are indistinguishable and thus we have decided to show only the ones corresponding to the former case. Thus (S.31) becomes:

$$\dot{\alpha}_{\sigma i} = \alpha_{\sigma i}\left[d\delta_\sigma \frac{\partial g_\sigma}{\partial \alpha_{\sigma i}} - \Theta(\varphi(\vec{\alpha}_\sigma))\frac{\partial\varphi(\vec{\alpha}_\sigma)/\partial\alpha_{\sigma i}}{\sum_{\tau=1}^m \sum_{k=1}^p \left(\partial\varphi(\vec{\alpha}_\sigma)/\partial\alpha_{\tau k}\right)^2 \alpha_{\tau k}} \sum_{\rho=1}^m \sum_{j=1}^p \frac{\partial\varphi(\vec{\alpha}_\sigma)}{\partial\alpha_{\rho j}} d\delta_\rho \alpha_{\rho j}\frac{\partial g_\sigma}{\partial\alpha_{\rho j}}\right] . \quad \text{(S.32)}$$

where we have also included the possibility that the constraint is taken into account as in (S.27).
As stated in the Main Text, we must now introduce a trade-off in the utilization of resources; we can do so by requiring that each species has a maximum total resource uptake rate, i.e. $\sum_{i=1}^p \alpha_{\sigma i} \leq E_\sigma^*$. If we now use $g_\sigma = \sum_{i=1}^p v_i \alpha_{\sigma i} r_i - \delta_\sigma$ and $\varphi(\vec{\alpha}_\sigma) = \sum_{i=1}^p \alpha_{\sigma i}/E_\sigma^* - 1$ (where we have rearranged the constraint to make it nondimensional) in (S.32), we obtain:

$$\dot{\alpha}_{\sigma i} = \alpha_{\sigma i} d\delta_\sigma \left[v_i r_i - \frac{\Theta\left(\varphi(\vec{\alpha}_\sigma)\right)}{\sum_{k=1}^p \alpha_{\sigma k}} \sum_{j=1}^p v_j r_j \alpha_{\sigma j}\right] . \quad \text{(S.33)}$$

In all the numerical calculations performed in this work, we approximated Heaviside's step function with the smooth sigmoid $\Theta(x) = 1/(1 + \exp(-x \cdot 10^{10}))$. While this function may seem very sharp, simulations performed with $\Theta$ defined as any non-negative and monotonously increasing function such that $\Theta(x) \approx 0$ for $x < 0$ and $\Theta(0) = 1$ (e.g. $\Theta(x) = \exp(k \cdot x)$ for many different $k > 1$, or $\Theta(x) = 1/[1 + \exp(-k \cdot (x + a))]$ with $k > 1$ and $a > 0$ chosen so that $\Theta$ satisfies the aforementioned properties) give very similar or totally indistinguishable outcomes.

#### B. Comparison between the model and experimental measures of diauxic growth curves

##### 1. Methods

*a. Experiment.* The *S. cerevisiae* strain used in this study, yAG47, is identical to strain yJHK459 of [1] and is in the W303 background. Its genotype is *MATa*, *can1-100*, *ura3*Δ0, *BUD4-S288C*. A culture of yAG47 was grown overnight in complete synthetic medium (CSM) + 2% (w/v) glucose. 1 mL of the overnight culture was spun down and resuspended in CSM + 0.5% (w/v) galactose to a concentration of $1.6 \cdot 10^5$ cells/mL. Eight wells of a 96-well plate were inoculated with 150 $\mu$L of the resuspended culture and incubated with constant shaking at 30°C in a plate reader. The 96-well plate was sealed with a sealing membrane that allowed gas exchange. The temperature on the top of the 96-well plate was kept at 31°C to avoid condensation on the membrane. Optical density (OD) measurements were taken every 10 min, for a total duration of about 70 h. To build the calibration curve used to convert OD to cell density, 1.4 mL of the same overnight culture were spun down and resuspended in 1 mL of CSM + 0.5% (w/v) galactose. The density of this suspension was measured using a Coulter counter and serial dilutions of this suspension were inoculated in a 96-well plate covered with the same sealing membrane used for the growth curve measurement. The OD of the wells containing the serial dilution of the suspension was measured after equilibration to 30°C using the same plate reader used to measure the growth curves, and these measurements were used to build the calibration curve converting OD to cell density.

*b. Model.* As explained in the Main Text, when *S. cerevisiae* is grown on galactose as the primary carbon source, the sugar is partially respired and partially fermented. As a byproduct of fermentation the yeast excretes ethanol, which can then be used as the second resource. For this reason, in order to describe such system we use our adaptive consumer-resource model with $m = 1$, $p = 2$ and we slightly modify the equations for the time evolution of ethanol concentration to take into account the fact that this resource is not initially present in the system but is produced by the yeast. In particular, the equations we use in order to describe the system are the following:

$$\dot{n} = n\left(v_{\text{gal}}\alpha_{\text{gal}}\frac{c_{\text{gal}}}{K_{\text{gal}} + c_{\text{gal}}} + v_{\text{eth}}\alpha_{\text{eth}}\frac{c_{\text{eth}}}{K_{\text{eth}} + c_{\text{eth}}} - \delta\right) , \quad \text{(S.34a)}$$

$$\dot{c}_{\text{gal}} = -n\alpha_{\text{gal}}\frac{c_{\text{gal}}}{K_{\text{gal}} + c_{\text{gal}}} , \quad \text{(S.34b)}$$

$$\dot{c}_{\text{eth}} = -n\alpha_{\text{eth}}\frac{c_{\text{eth}}}{K_{\text{eth}} + c_{\text{eth}}} + Y \cdot n\alpha_{\text{gal}}\frac{c_{\text{gal}}}{K_{\text{gal}} + c_{\text{gal}}} , \quad \text{(S.34c)}$$

$$\dot{\alpha}_i = \alpha_i d\delta \left[ v_i \frac{c_i}{K_i + c_i} - \Theta\left( \frac{\alpha_{\text{gal}} + \alpha_{\text{eth}}}{\mathcal{Q}\delta} - 1 \right) \frac{1}{\alpha_{\text{gal}} + \alpha_{\text{eth}}} \cdot \left( v_{\text{gal}} \alpha_{\text{gal}} \frac{c_{\text{gal}}}{K_{\text{gal}} + c_{\text{gal}}} + v_{\text{eth}} \alpha_{\text{eth}} \frac{c_{\text{eth}}}{K_{\text{eth}} + c_{\text{eth}}} \right) \right] \qquad i = \text{gal, eth} \tag{S.34d}$$

$$\frac{\alpha_{\text{gal}} + \alpha_{\text{eth}}}{\mathcal{Q}\delta} \leq 1 \ , \tag{S.34e}$$

where in (S.34c) we have inserted an ethanol production rate that is proportional to the galactose consumption rate; in other words, $Y$ can be interpreted as the galactose-to-ethanol yield.

##### *2. Results*

We have then performed a fit of the consumer-resource model to the experimental measure of the population density of *S. cerevisiae*, both in the case of adaptive and fixed metabolic strategies. The comparison between the data and the best-fits of the two models are shown in Figure 1 of the Main text, while the values of the parameters obtained are shown in Table S.1. As we can see from Figure 1 of the Main Text, the model with adaptive metabolic strategies can reproduce the experimental data, while the classic consumer-resource model with fixed $\alpha_{\sigma i}$-s cannot.
Notice that since the model has several parameters (10 in total, some of which are phenomenological) there can be several different choices that lead to apparently equivalent fits. The results that we reported in this work are those, among the one we have explored, that give the lowest absolute value of the Akaike Information Criterion (AIC) [2]. In particular, when fitting the model (both in the case of adaptive and fixed metabolic strategies) to the data we have found that the lowest AIC value was obtained when $\alpha_{\text{gal}}(0)$ and $\alpha_{\text{eth}}(0)$ were fixed and all other parameters were used as fitting parameters. We have therefore varied $\alpha_{\text{gal}}(0)$ and $\alpha_{\text{eth}}(0)$ along a grid and selected the values that minimize the AIC.
Once the fits have been determined, we have found that the adaptive model gives a much better fit to the data as measured by the AIC. In fact, if we call $\Delta_{\text{AIC}} := \text{AIC}_{\text{adaptive}} - \text{AIC}_{\text{fixed}}$ the difference between the AIC of the two models, it is possible to show [2, Sections 2.6-2.9] that $\exp(\Delta_{\text{AIC}}/2)$ is the relative likelihood of the two models, and as such measures the probability that the model with fixed metabolic strategies minimizes the information loss (i.e. it is a better fit to the data than the one with adaptive metabolic strategies). In our case, we found $\Delta_{\text{AIC}} = -2739.5$, so the probability that the results could be better explained using fixed metabolic strategies is infinitesimal.
Finally, in Figure S.2 we show how the model reproduces the evolution in time of the resources concentrations, metabolic strategies and also of the constraint (S.34e) using the parameters shown in Table S.1.

#### III. COEXISTENCE OF MULTIPLE SPECIES

##### A. Extinction times

As shown in I A, if the $E^*_\sigma$s are completely independent parameters, i.e. $E^*_\sigma/\delta_\sigma = \mathcal{Q}_\sigma$, we cannot find a feasible solution of (S.2b) and so the system obeys the CEP, i.e. no more than $p$ species will be able to coexist. However, as Figure 2 of the Main Text shows, the length of the time interval during which the species are able to coexist increases as the CTRs $\mathcal{Q}_\sigma = E^*_\sigma/\delta_\sigma$ are increasingly peaked around their mean value $\langle \mathcal{Q} \rangle$. This suggests that coexistence should be possible if each species was characterized by a unique timescale, i.e. if $E^*_\sigma/\delta_\sigma = \mathcal{Q}\ \forall\sigma$ (and our discussion in section I A shows that this is indeed the case), which is an assumption in agreement with the metabolic theory of ecology.

The extinction times shown in Fig. 2 have been computed as follows: we have drawn $\mathcal{Q}_\sigma \in \mathcal{N}[10^-6, \Sigma]$ g of resource/cells (with $\mathcal{N}$ the normal distribution), $\delta_\sigma \in \mathcal{U}[5 \cdot 10^{-3}, 5 \cdot 10^{-2}]$ 1/h, $E_\sigma(0) \in \mathcal{U}[0, \mathcal{Q}_\sigma\delta_\sigma]$, $v_i \in \mathcal{U}[10^8, 5 \cdot 10^9]$ cells/g of resource, $n_\sigma(0) \in \mathcal{U}[10^6, 5 \cdot 10^6]$ cells/mL, $c_i(0) \in \mathcal{U}[10^{-3}, 10^{-2}]$ g of resource/mL, $K_i \in \mathcal{U}[10^{-4}, 10^{-3}]$ g of resource/mL. Then, for each iteration we have drawn $s_i \in \mathcal{U}[10^{-3}, 10^{-2}]$ g of resource/(mL·h) and $\alpha_{\sigma i}(0)$ so that $\sum_{i=1}^{p} \alpha_{\sigma i}(0) = E_\sigma(0)$, simulated the equations and computed the instant at which for the first, third, fifth and seventh time population of one species dropped below 1 cell/mL.

##### B. Metabolic theory of ecology

Here, we explore this point in more detail. To interpret our model in view of the metabolic theory of ecology, we need to vary slightly the notation and rewrite Eqs. 1-2 of the main text (or Eqs. (S.1a-S.1b) above) replacing the population density $n_\sigma$ with the total biomass density, $M_\sigma$, of species $\sigma$. If $\mathcal{M}_\sigma$ is the characteristic mass of an individual of species $\sigma$, the two variables are linked by $M_\sigma = \mathcal{M}_\sigma n_\sigma$. Eqs. 1-2 in this framework read:

$$\dot{M}_\sigma = M_\sigma \left( \sum_{i=1}^{p} v_i \alpha_{\sigma i} r_i(c_i) - \delta_\sigma \right) , \tag{S.35}$$

$$\dot{c}_i = s_i - \sum_{\sigma=1}^{m} M_\sigma \alpha_{\sigma i} r_i(c_i) - \mu_i c_i , \tag{S.36}$$

where $\alpha_{\sigma i}$ is the maximum uptake rate of resource $i$ by species $\sigma$, per unit mass of species $\sigma$. Note that even though Eqs. (S.35-S.36) are identical to s. (S.1a-S.1b) after replacing $n_\sigma$ with $M_\sigma$, they are not exactly equivalent because (S.36) and (S.1b) of the Main Text differ by a factor $\mathcal{M}_\sigma$ within the summation. This difference does not affect the main results of our work, and we decided to present our results in the main text in terms of $n_\sigma$ rather than $M_\sigma$ for consistency with previous works. The metabolic theory of ecology prescribes that

$$\alpha_{\sigma i} \propto \mathcal{M}_\sigma^{-\lambda} \qquad \text{and} \qquad \delta_\sigma \propto \mathcal{M}_\sigma^{-\lambda} \tag{S.37}$$

with $\lambda$ a universal exponent. Brown et al. (*Ecology*, 2004) have suggested $\lambda = 1/4$, but our results are independent on its particular value. Since $E^*_\sigma = \sum_{i=1}^{p} \alpha_{\sigma i}(t)$ we have $E^*_\sigma \propto \mathcal{M}_\sigma^{-\lambda}$, and thus the ratio $E^*_\sigma/\delta_\sigma = \mathcal{Q}$ is independent of $\sigma$, which is the condition for which coexistence beyond the CEP is possible in our framework. In other words, coexistence in our framework is possible if each species is characterized by a unique timescale that is proportional to $\mathcal{M}_\sigma^{-\lambda}$.

##### C. Unfavorable resources

As we have shown in I A, when using the soft constraint $\sum_{i=1}^{p} \alpha_{\sigma i} \leq \mathcal{Q}\delta_\sigma$ and (S.33) we must have $1/v_i < \mathcal{Q}\ \forall i$; the system, however, behaves interestingly also when we have $1/v_j > \mathcal{Q}$ for some $j$. In Figure S.4 we show the outcome of a numerical integration of the system in this case; as we can see, what happens is that the components of the metabolic strategies relative to the unfavorable resource (i.e. the one for which we have $1/v_j > \mathcal{Q}$) go to zero, so the species are stopping wasting energy for the metabolization of such resource.
Something analogous happens also when the degradation rates $\mu_i$ are non-null in (S.1b); in particular if one of the resources is sufficiently volatile, i.e. $\mu_j$ is sufficiently larger than $1/v_j$, all the $j$-th components of the metabolic strategies will vanish. Figure S.5 shows an example of this case, while Figure S.7 shows how increasing the magnitude of the degradation rates makes the various components of $\vec{\alpha}_\sigma$ gradually vanish.

##### D. Variable environmental conditions

As stated in the Main Text, using adaptive metabolic strategies has positive effects when the nutrient supply rate vector changes with time. To study how the system behaves in this case, we have used the following rectangular wave temporal dynamics for $\vec{s}(t)$:

$$\vec{s}(t) = \begin{cases} \vec{s}_{\text{in}} & 0 < t' \leq \tau_{\text{in}} \\ \vec{s}_{\text{out}} & \tau_{\text{in}} < t' \leq \tau_{\text{out}} \end{cases} , \tag{S.38}$$

where $\vec{s}_{\text{in}}$ and $\vec{s}_{\text{out}}$ are two fixed values (i.e., set at the start of the simulations) chosen such that their rescaled versions $\hat{\vec{s}}_{\text{in}}$ and $\hat{\vec{s}}_{\text{out}}$ are, respectively, inside and outside of the convex hull of the initial metabolic strategies, and $t' = t - \tau\lfloor t/\tau \rfloor$ with $\tau = \tau_{\text{in}} + \tau_{\text{out}}$.

We show in Figures S.6 the temporal dynamics of all time-varying quantities for the simulation shown in Figure 4 of the Main Text, where $\vec{s}(t)$ is given by (S.38); as shown in the figure, when metabolic strategies evolve with time, the populations have increased mean densities and reduced oscillations, compared to the case of fixed metabolic strategies. Furthermore, when the $\vec{\alpha}_\sigma$s are fixed, the difference between $\tau_{\text{in}}$ and $\tau_{\text{out}}$ is crucial to determine if the CEP holds or not: we show in Figures S.6b and S.6c the temporal dynamics of all time-varying quantities for the same system shown in Figures S.6d and S.6e, but with a smaller value for $\tau_{\text{in}}$. In this case, we can see that all species manage to coexist even with fixed metabolic strategies.

Finally, if we also let $\mu_i \neq 0$ we recover something similar to what we have shown in III C: while the metabolic strategies continue to oscillate, if $\mu_j$ is sufficiently large then all the $j$-th components of $\vec{\alpha}_\sigma$ will vanish, as shown in Figure S.8.

##### E. Slow adaptation can still lead to competitive exclusion

From what we have stated in the Main Text and shown in I A, it could seem that our model introduces yet another paradox because it predicts that with adaptive metabolic strategies the CEP is *always* violated; however, from experiments we know that, for example, competition between two microbial species can either result in exclusion or coexistence (Friedman et al., *Nature Ecology and Evolution*, 2017). The key aspect of our model that determines the actual outcome of competition is the adaptation velocity $d$: if it is sufficiently small, in fact, competitive exclusion can happen. As Figure S.9 shows, as $d \to 0$, the stationary populations $n_\sigma^*$ of the species with higher ranks become smaller and smaller by orders of magnitude, up to the point where most of the species go extinct. On the other hand, if $\vec{s}(t)$ changes with time, then fast adaptation (i.e. large $d$) will result in less variable populations; in particular, once the parameters and initial conditions have been fixed and the equations solved, for each solution $n_\sigma(t)$ we have computed the mean population as

$$\langle n_\sigma \rangle = \frac{1}{T}\int_0^T n_\sigma(t)dt \ , \tag{S.39}$$

where $T$ is the length of the integration interval, and then we have computed the "variance" of $n_\sigma(t)$ as

$$\Sigma^2_{n_\sigma} = \frac{1}{T}\int_0^T (n_\sigma(t) - \langle n_\sigma \rangle)^2 dt \tag{S.40}$$

and studied the distribution of the coefficients of variation $\Sigma_{n_\sigma} / \langle n_\sigma \rangle$ as $d$ increases. The results are shown in Figure S.10: as we can see, as $d$ grows the distribution of $\Sigma_{n_\sigma} / \langle n_\sigma \rangle$ is increasingly peaked around small values, meaning exactly that as adaptation velocity increases the species' population become more stable.

---

[1] J. H. Koschwanez, K. R. Foster, and A. W. Murray, eLife **2**, e00367 (2013).
[2] K. P. Burnham and D. R. Anderson, *Model Selection and Multimodel Inference: A practical information-theoretic approach*, 2nd ed. (Springer Verlag, 2002).
[3] P. Malakar and K. V. Venkatesh, FEMS Yeast Research **14**, 346 (2014).
[4] B. Sonnleitner and O. Käppeli, Biotechnology and Bioengineering **28**, 927 (1986).
[5] J. H. Kim, J. Ryu, I. Y. Huh, S.-K. Hong, H. A. Kang, and Y. K. Chang, Bioprocess and Biosystems Engineering **37**, 1871 (2014).
[6] T. Vos, X. D. V. Hakkaart, E. A. F. de Hulster, A. J. A. van Maris, J. T. Pronk, and P. Daran-Lapujade, Microbial Cell Factories **15**, 111 (2016).
[7] C. Bro, S. Knudsen, B. Regenberg, L. Olsson, and J. Nielsen, Applied and Environmental Microbiology **71**, 6465 (2005).
[8] P. Dantigny, Journal of Biotechnology **43**, 213 (1995).

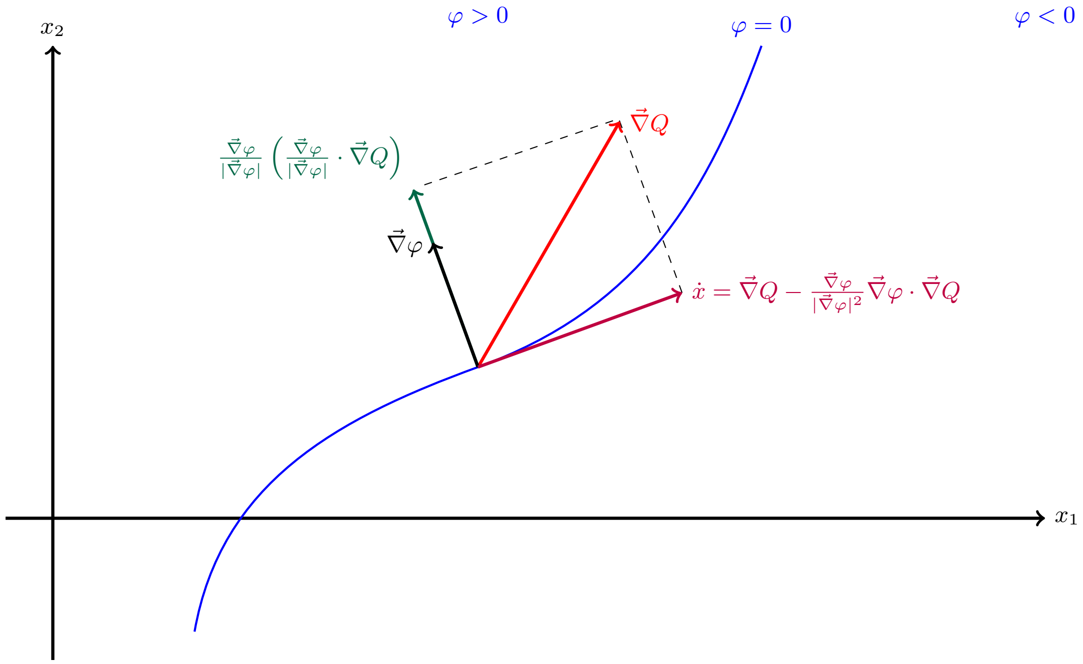


**FIG. S.1:** Effect of the additional term introduced in (S.24). This is just a descriptive (and not realistic) example made for the case $p = 2$ for ease of representation, to make it easier to understand that the additional term in (S.24) bounds $\vec{x}(t)$ to move on the manifold defined by the constraint $\varphi(\vec{x}) = 0$.

**TABLE S.1:** Values of the parameters of the model obtained from the best fit, both in the case with adaptive and fixed metabolic strategies. The fit was performed with Mathematica's `NonlinearModelFit` function, and the results have been approximated to the third significant figure.

| **Parameter** | **Value (adaptive $\alpha_{\sigma i}$)** | **Value (fixed $\alpha_{\sigma i}$)** | **Value from the literature** | **Units** |
|---|---|---|---|---|
| $v_{\rm gal}$ | $1.20 \cdot 10^{10}$ | $1.08 \cdot 10^{10}$ | N/A[a] | cells/g of resource |
| $v_{\rm eth}$ | $1.28 \cdot 10^{10}$ | $2.00 \cdot 10^{10}$ | N/A[a] | cells/g of resource |
| $K_{\rm gal}$ | $1.41 \cdot 10^{-3}$ | $2.23 \cdot 10^{-9}$ | $1.7 \cdot 10^{-4}$ [3][b] | g of resource/mL |
| $K_{\rm eth}$ | $9.81 \cdot 10^{-3}$ | $2.24 \cdot 10^{-2}$ | $10^{-4}$ [4, Table I][b] | g of resource/mL |
| $Y$ | 0.52 | 0.45 | 0.41 [5, Table 1][c] | g of ethanol/g of galactose |
| $\mathcal{Q}$ | $7.63 \cdot 10^{-6}$ | N/A | N/A | g of resource/cells |
| $\delta$ | $6.14 \cdot 10^{-6}$ | $2.56 \cdot 10^{-3}$ | $4.7 \cdot 10^{-4}$ [6][d] | 1/h |
| $\alpha_{\rm gal}(0)$[e] | $1.44 \cdot 10^{-11}$ | $2.49 \cdot 10^{-11}$ | $3.60 \cdot 10^{-11}$[7, Table 3][f] | g of resource/(cells · h) |
| $\alpha_{\rm eth}(0)$[e] | $7.74 \cdot 10^{-12}$ | $1.65 \cdot 10^{-11}$ | N/A | g of resource/(cells · h) |
| $d$ | $1.47 \cdot 10^{-6}$ | N/A | N/A[a] | (nondimensional) |

[a] This is a phenomenological parameter that cannot be directly measured.
[b] As stated in [8], the values of the half-saturation constants can change sensibly depending on the medium used in the experiment. It is reasonable to expect that at high concentrations of galactose or ethanol, other nutrients in the medium become the limiting resource. As a consequence, the Monod half-saturation constant is not a property of the carbon source by itself but depends on the whole set of nutrients available to the cell. Thus, it is not surprising finding a discrepancy between the estimate of our fits and measures taken from the literature coming from experiments done in very different conditions.
[c] We have performed an average of the measures reported in the cited table. Notice that the experiment in [5] was performed with initial concentrations of galactose different from the one we used.
[d] The cited experiment was performed in different conditions with respect to our experiment. Notice also that our fits may provide an underestimation of $\delta$, since dead cells could still contribute to the OD of a culture.
[e] This parameter has been fixed, and it was not used as a free parameter for the fit (see section II B 2).
[f] This value refers to the maximum galactose uptake rate; compare it to the evolution of the galactose metabolic strategy in Figure S.2b.

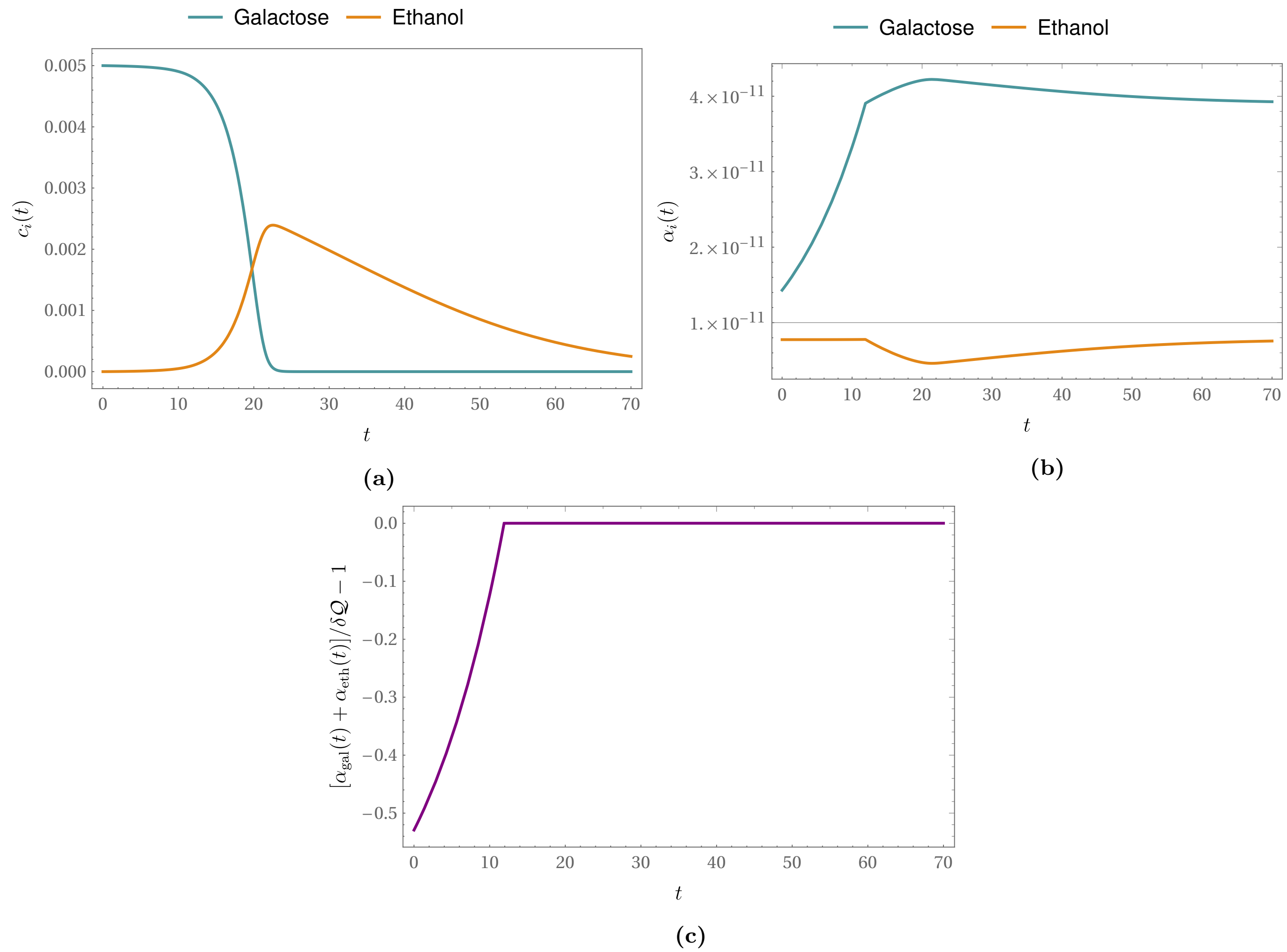


**FIG. S.2:** Time evolution of the consumer-resource model with adaptive metabolic strategies when using the parameters found by our fit, as shown in Table S.1. The time evolution of the population is of course given in Figure 1A of the Main Text. **(a)**: Time evolution of the resources' concentrations. Notice that as galactose is consumed ethanol is produced, and then consumed only when galactose is depleted. **(b)**: Time evolution of the metabolic strategies. Notice that when the upper bound on the total uptake rate is reached, the increase in $\alpha_{\rm gal}$ is balanced by a reduction of $\alpha_{\rm eth}$, and when galactose is depleted $\alpha_{\rm eth}$ starts growing. **(c)**: Time evolution of the constraint (S.34e) as rewritten in (S.33).

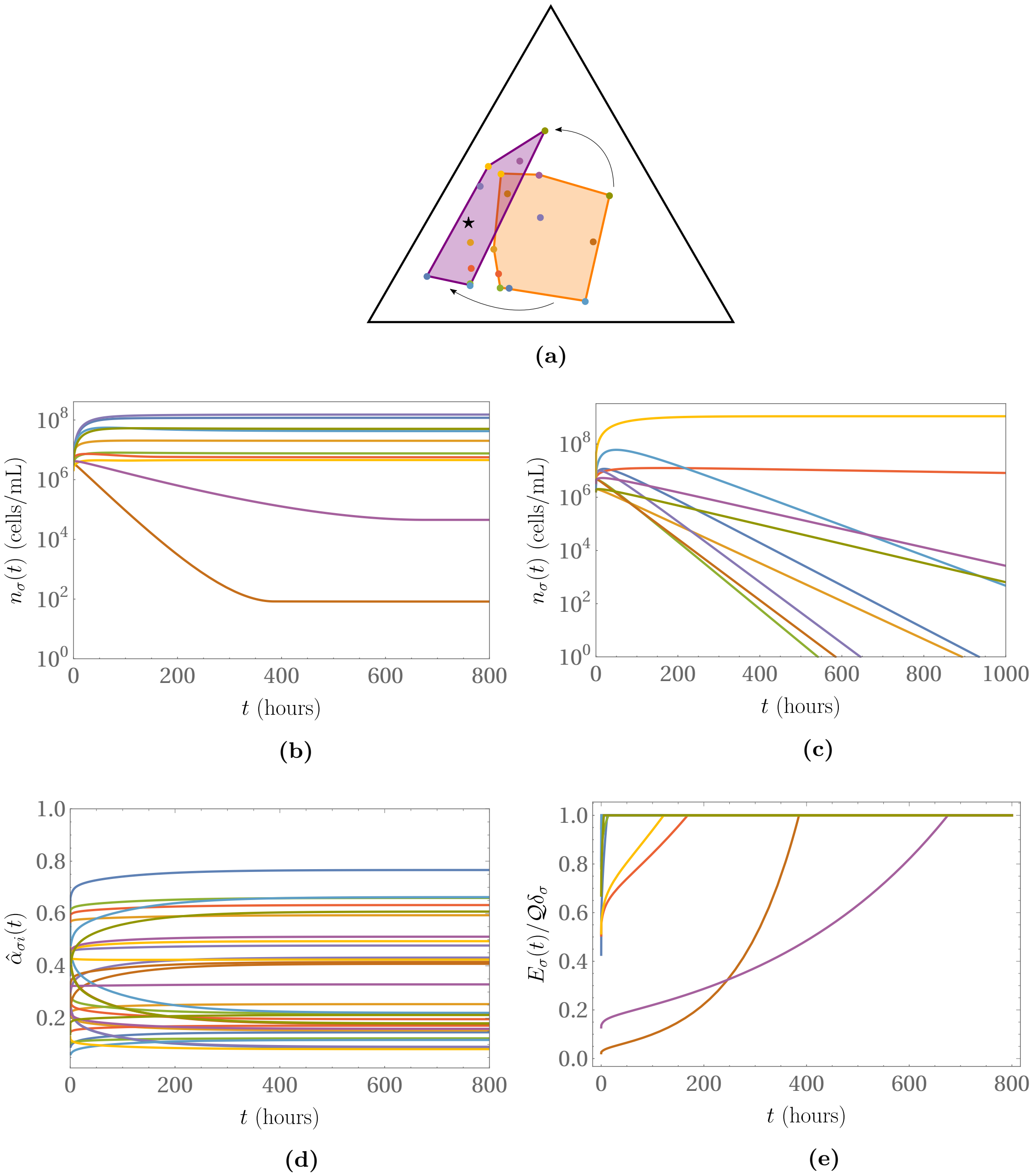


**FIG. S.3:** Comparison between the time evolution of a "classic" consumer-resource model (i.e. with fixed metabolic strategies) and our adaptive version with (S.33). In this simulation we have considered a system with $m = 10$, $p = 3$, $\mathcal{Q} \in \mathcal{U}[10^{-7}, 5 \cdot 10^{-5}]$ g of resource/cells (with $\mathcal{U}$ the uniform distribution), $\delta_\sigma \in \mathcal{U}[5 \cdot 10^{-3}, 5 \cdot 10^{-2}]$ 1/h, $E_\sigma(0) \in \mathcal{U}[0, \mathcal{Q}\delta_\sigma]$ g of resource/(cells · h), $v_i \in \mathcal{U}[10^8, 5 \cdot 10^9]$ cells/g of resource, $n_\sigma(0) \in \mathcal{U}[10^6, 5 \cdot 10^6]$ cells/mL, $c_i(0) \in \mathcal{U}[10^{-3}, 10^{-2}]$ g of resource/mL, $K_i \in \mathcal{U}[10^{-4}, 10^{-3}]$ g of resource/mL. We have drawn $s_i \in \mathcal{U}[10^{-3}, 10^{-2}]$ g of resource/(mL·h) and $\alpha_{\sigma i}(0)$ $\sum_{i=1}^{p} \alpha_{\sigma i}(0) = E_\sigma(0)$. **(a)**: Comparison between the initial (orange) and final (purple) convex hull of the rescaled metabolic strategies $\hat{\alpha}_{\sigma i}$ (colored dots) when they are allowed to evolve (rescaled quantities have been defined in (S.8)). As we can see, in the final state $\hat{\alpha}_{\sigma i}$ have "incorporated" the rescaled nutrient supply rate vector $\vec{s}$ (black star) in their convex hull. The triangle is the 2-dimensional simplex on which both $\vec{s}$ and $\vec{\alpha}_\sigma$ lie. See the SI webpage for the animation of the time evolution of the convex hull. **(b) and (c)**: Comparison of the time evolution of the species' populations between the case of adaptive (b) and static (c) metabolic strategies, with the *same* initial conditions (the colors of the curves match the corresponding $\hat{\alpha}_{\sigma i}$ in (a)). **(d)**: Time evolution of the rescaled metabolic strategies (each curve represent one of the components of $\vec{\alpha}_\sigma$, and the colors match the strategies in (a)). **(e)**: Time evolution of the ratios $E_\sigma(t)/\mathcal{Q}\delta_\sigma$; as we can see, they all saturate to their maximum value allowed; the colors of the curves match the corresponding strategy in (a).

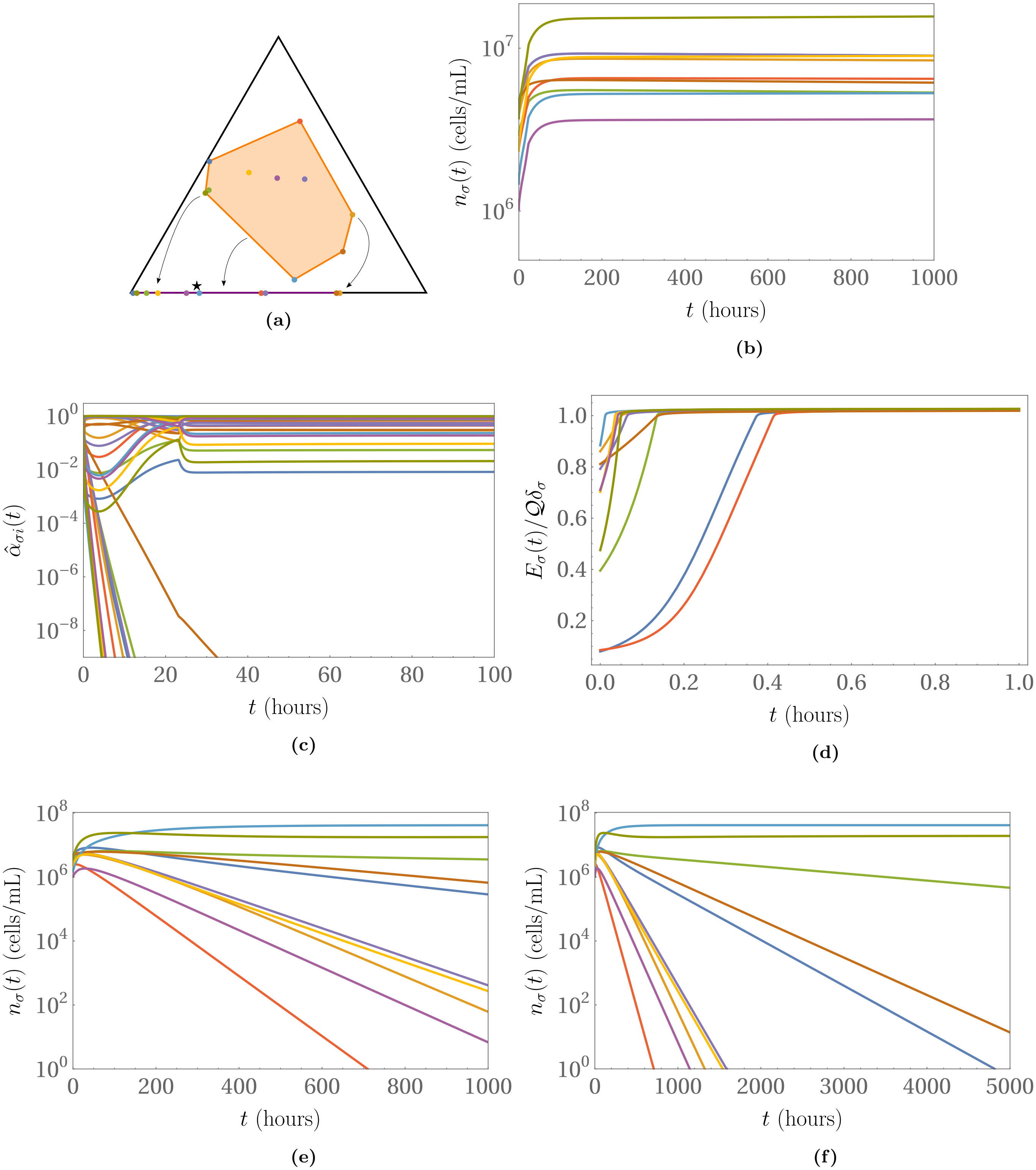


**FIG. S.4:** Another instance of integration of the model with (S.33) for the time evolution of the metabolic strategies. We have used $m = 10$, $p = 3$, $\mathcal{Q} \in \mathcal{U}[10^{-8}, 10^{-7}]$ g of resource/cells, $\delta_\sigma \in \mathcal{U}[5 \cdot 10^{-3}, 5 \cdot 10^{-2}]$ 1/h, $E_\sigma(0) \in \mathcal{U}[0, \mathcal{Q}\delta_\sigma]$, $n_\sigma(0) \in \mathcal{U}[10^6, 5 \cdot 10^6]$ cells/mL, $c_i(0) \in \mathcal{U}[10^{-3}, 10^{-2}]$ g of resouce/mL, $K_i \in \mathcal{U}[10^{-4}, 10^{-3}]$ g of resource/mL; we have then set $\vec{v} = (9 \cdot 10^6, 5.45 \cdot 10^8, 6.5 \cdot 10^7)$, so that $1/v_1 = 1.11 \cdot 10^{-7} > \mathcal{Q}$ and $1/v_2$, $1/v_3 < \mathcal{Q}$. Furthermore, we have drawn $s_i \in \mathcal{U}[10^-3, 10^-2]$ g of resource/(mL·h) and $\alpha_{\sigma i}(0)$ $\sum_{i=1}^{p} \alpha_{\sigma i}(0) = E_\sigma(0)$. In order to be represented on the same simplex, the metabolic strategies and the nutrient supply rate vector have been rescaled as usual. **(a)**: Comparison between the initial (orange) and final (purple) convex hull of the metabolic strategies (colored dots). As we can see, in the final state all metabolic strategies have "squeezed" onto the same side of the simplex. See the SI webpage for the animation of the time evolution of the system. **(b)**: Time evolution of the species' populations. **(c)**: Time evolution of the rescaled metabolic strategies; as we can see, some of their components indeed decay towards zero. **(d)**: Time evolution of the values of the ratios $E_\sigma(t)/\mathcal{Q}\delta_\sigma$. **(e) and (f)**: Time evolution of the same system with fixed metabolic strategies, where $E_\sigma = \mathcal{Q}\delta_\sigma$.

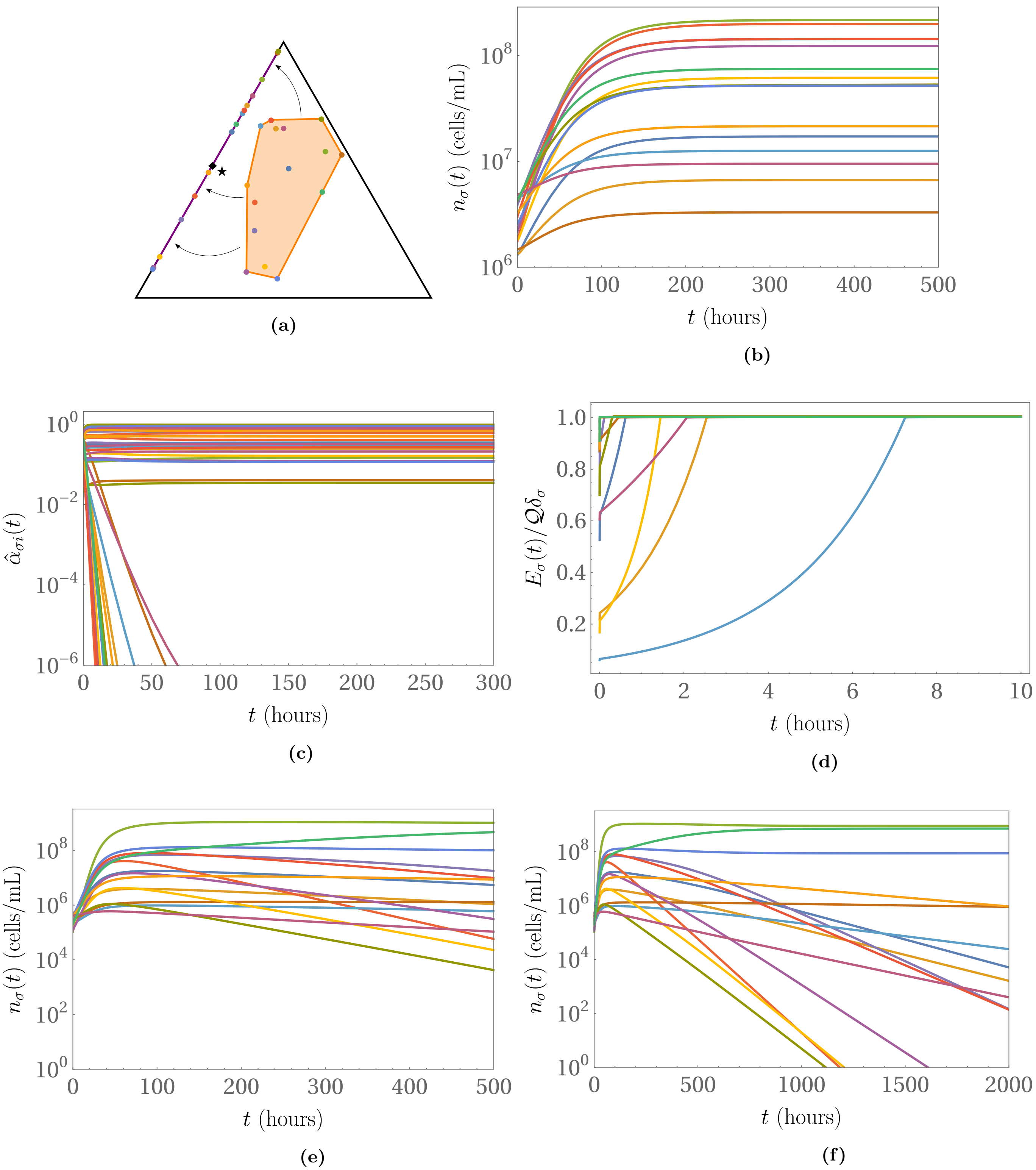


**FIG. S.5:** Integration of the model with (S.33) for the time evolution of the metabolic strategies and non-null degradation rates. We have used $m = 15$, $p = 3$, $\mathcal{Q} \in \mathcal{U}[10^{-7}, 5 \cdot 10^{-7}]$ g of resource/cells, $\delta_\sigma \in \mathcal{U}[5 \cdot 10^{-3}, 5 \cdot 10^{-2}]$ 1/h, $E_\sigma(0) \in \mathcal{U}[0, \mathcal{Q}\delta_\sigma]$, $v_i \in \mathcal{U}[10^9, 5 \cdot 10^9]$ cells/g of resource, $n_\sigma(0) \in \mathcal{U}[10^6, 5 \cdot 10^6]$, $c_i(0) \in \mathcal{U}[10^{-3}, 10^{-2}]$ g of resource/mL, $K_i \in \mathcal{U}[10^{-4}, 10^{-3}]$ g of resource/mL, $s_i \in \mathcal{U}[10^{-3}, 10^{-2}]$ g of resource/(mL·h) and $\mu_i \in \mathcal{U}[10^3, 10^4]$ 1/h. In order to be represented on the same simplex, the metabolic strategies and the nutrient supply rate vector have been rescaled as usual. **(a)**: Comparison between the initial (orange) and final (purple) convex hull of the metabolic strategies (colored dots). As we can see, $\vec{s}$ (black diamond) (see (S.15)) lies on one side of the simplex, so in the final state all metabolic strategies have squeezed onto that same side. See the SI webpage for the animation of the time evolution of the system. **(b)**: Time evolution of the species' populations. **(c)**: Time evolution of the rescaled metabolic strategies; as we can see, some of their components indeed decay towards zero. **(d)**: Time evolution of the values of the ratios $E_\sigma(t)/\mathcal{Q}\delta_\sigma$. **(e) and (f)**: Time evolution of the same system with fixed metabolic strategies, where $E_\sigma = \mathcal{Q}\delta_\sigma$.

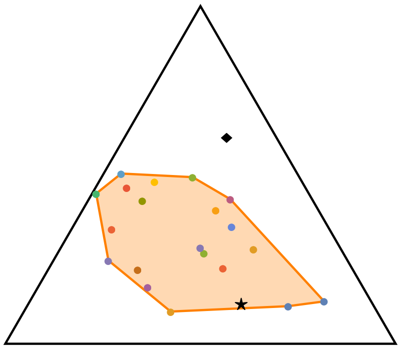

**(a)**

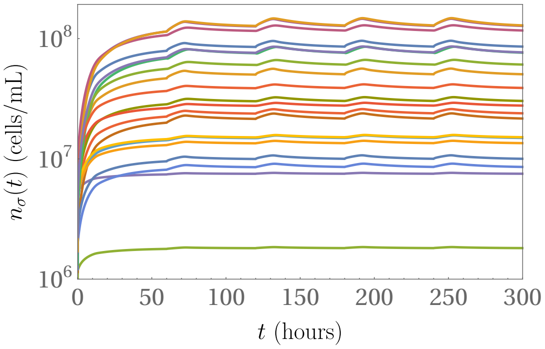


**(b)**

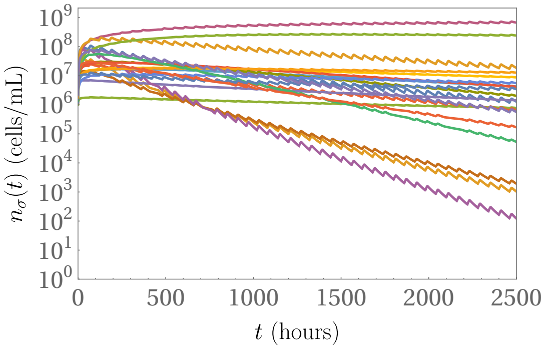


**(c)**

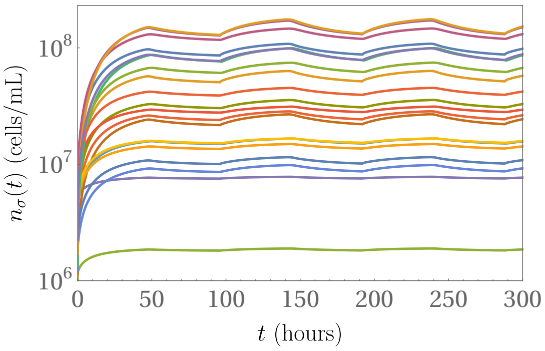


**(d)**

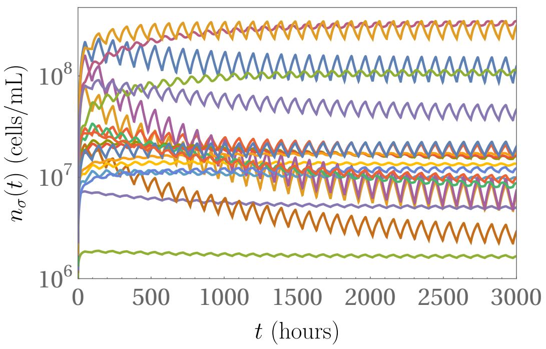


**(e)**

**FIG. S.6:** Comparison between the behavior of the classic MacArthur's model and our adaptive version, when $\vec{s}(t)$ is a function of time. We have used $m = 20$, $p = 3$, $\mathcal{Q} \in \mathcal{U}[10^{-7}, 5\cdot 10^{-6}]$ g of resource/cells, $\delta_\sigma \in \mathcal{U}[5\cdot 10^{-3}, 5\cdot 10^{-2}]$ 1/h, $E_\sigma(0) = \mathcal{Q}\delta_\sigma$ (so that the CEP can be violated in principle also when using fixed strategies), $v_i \in \mathcal{U}[10^8, 5\cdot 10^9]$ cells/g of resource, $n_\sigma(0) \in \mathcal{U}[10^6, 5\cdot 10^6]$ cells/mL, $c_i(0) \in \mathcal{U}[10^-3, 10^-2]$ g of resource/mL, $K_i \in \mathcal{U}[10^-4, 10^-3]$ g of resource/mL; $\vec{s}(t)$ is given as in (S.38), with the components of $\vec{s}_{\text{in}}$ drawn in $\mathcal{U}[10^-3, 10^-2]$ g of resource/(mL·h), so that its rescaled version lies in the convex hull of $\vec{\alpha}_\sigma$, and similarly for $\vec{s}_{\text{out}}$ (with its rescaled version falling outside of the convex hull). In order to be represented on the same simplex, the metabolic strategies and the nutrient supply rate vector have been rescaled as usual. Figure 4 of the Main Text contains the results shown here only for the population of species $\sigma = 20$. **(a)**: Initial configuration of the system, where both $\vec{s}_{\text{in}}$ (black star) and $\vec{s}_{\text{out}}$ (black diamond) are represented. See the SI webpage for the animation of the time evolution of the system. **(b)**: Time evolution of the species' populations using adaptive metabolic strategies and $\tau_{\text{in}} = 12$h and $\tau_{\text{out}} = 48$h. **(c)**: Time evolution of the species' populations using fixed metabolic strategies and $\tau_{\text{in}} = 12$h, $\tau_{\text{out}} = 48$h. **(d)**: Time evolution of the species' populations using adaptive metabolic strategies and $\tau_{\text{in}} = \tau_{\text{out}} = 48$h. **(e)**: Time evolution of the species' populations using fixed metabolic strategies and $\tau_{\text{in}} = \tau_{\text{out}} = 48$h.

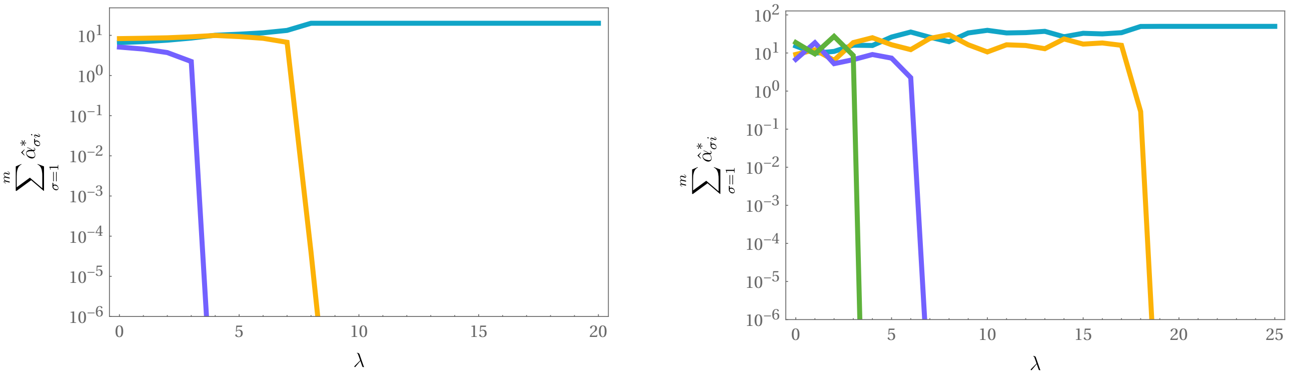


**(a)** **(b)**

**FIG. S.7:** Here we show what happens when the magnitude of the degradation rates is increased; in particular, we have substituted $-\mu_i c_i$ in (S.1b) with $-\lambda\mu_i c_i$, where $\lambda$ now regulates the magnitude of the degradation rates. We have used the following parameters: $\mathcal{Q} \in \mathcal{U}[10^{-7}, 10^{-6}]$ g of resource/cells, $\delta_\sigma \in \mathcal{U}[5 \cdot 10^{-3}, 5 \cdot 10^{-2}]$ 1/h, $E_\sigma(0) \in \mathcal{U}[0, \mathcal{Q}\delta_\sigma]$, $v_i \in \mathcal{U}[10^8, 5 \cdot 10^9]$ cells/g of resource, $n_\sigma(0) \in \mathcal{U}[10^6, 5{\cdot}10^6]$ cells/mL, $c_i(0) \in \mathcal{U}[10^{-3}, 10^{-2}]$ g of resource/mL, $K_i \in \mathcal{U}[10^{-4}, 10^{-3}]$ g of resource/mL, $\mu_i \in \mathcal{U}[10^3, 5 \cdot 10^3]$ 1/h; $\alpha_{\sigma i}(0)$ have been drawn so that $\sum_{i=1}^{p} \alpha_{\sigma i}(0) = E_\sigma(0)$, and $s_i$ have been drawn in $\mathcal{U}[10^{-3}, 10^{-2}]$ g of resource/(mL·h) so that $\vec{\hat{s}}$ belongs to the convex hull of the rescaled metabolic strategies. Furthermore, we have ordered $v_i$ and $\mu_i$ so that $1/v_i < \cdots < 1/v_p$ and $\mu_1 < \cdots < \mu_p$. Using always the same parameters and initial conditions we have computed the value of $\sum_{\sigma=1}^{m} \hat{\alpha}^*_{\sigma i}$, i.e. the total uptake rate of each resource $i$, for increasing values of $\lambda$. As we can see, in both (a) and (b) we have that when $\lambda$ is sufficiently large the $p$–th component of $\sum_{\sigma=1}^{m} \hat{\alpha}^*_{\sigma i}$ vanishes, i.e. the species have stopped using resource $p$; then, as $\lambda$ increases, the same happens for resource $p-1$, $p-2$, and so on until only resource 1 is used. **(a)**: In this case we have used $m = 20$ and $p = 3$. **(b)**: In this case we have used $m = 50$ and $p = 4$.

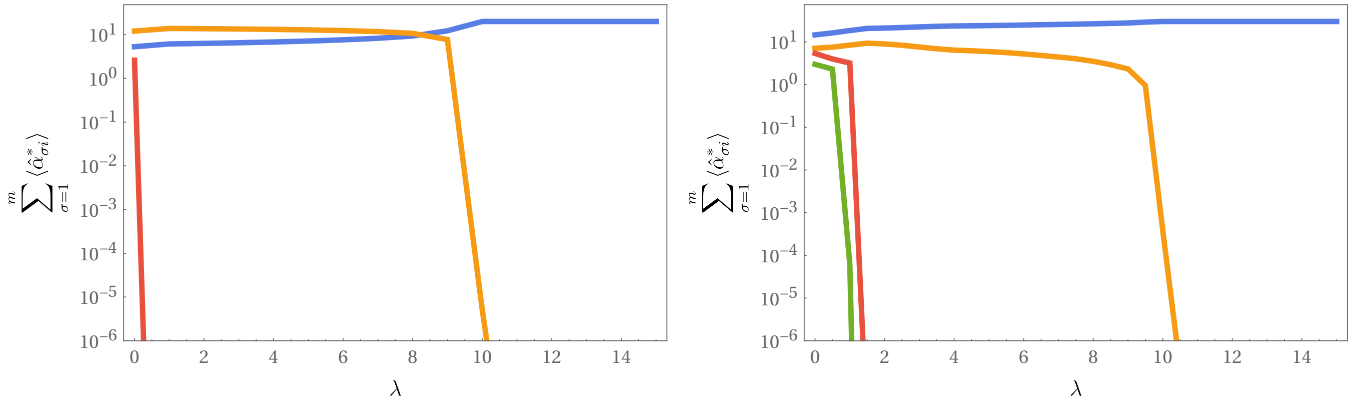


**(a)** **(b)**

**FIG. S.8:** Here we show what happens when $\vec{s}$ changes in time and the magnitude of the degradation rates is increased. We have drawn the parameters from the same distributions of Figure S.7, but this time $\vec{s}(t)$ is given by (S.38). Furthermore, in this case we have computed the sum $\sum_{\sigma=1}^{m} \langle \hat{\alpha}_{\sigma i} \rangle$ using the average value of the metabolic strategies, computed as $\langle \hat{\alpha}_{\sigma i} \rangle = \langle \alpha_{\sigma i} \rangle / \sum_{j=1}^{p} \langle \alpha_{\sigma i} \rangle$, with $\langle \alpha_{\sigma i} \rangle = \frac{1}{T/2} \int_{T/2}^{T} \alpha_{\sigma i}(t)dt$. **(a)**: In this case we have used $m = 20$, $p = 3$, $\tau_{\text{in}} = 48$ h and $\tau_{\text{out}} = 24$ h. **(b)**: In this case we have used $m = 30$, $p = 4$ and $\tau_{\text{in}} = \tau_{\text{out}} = 36$ h.

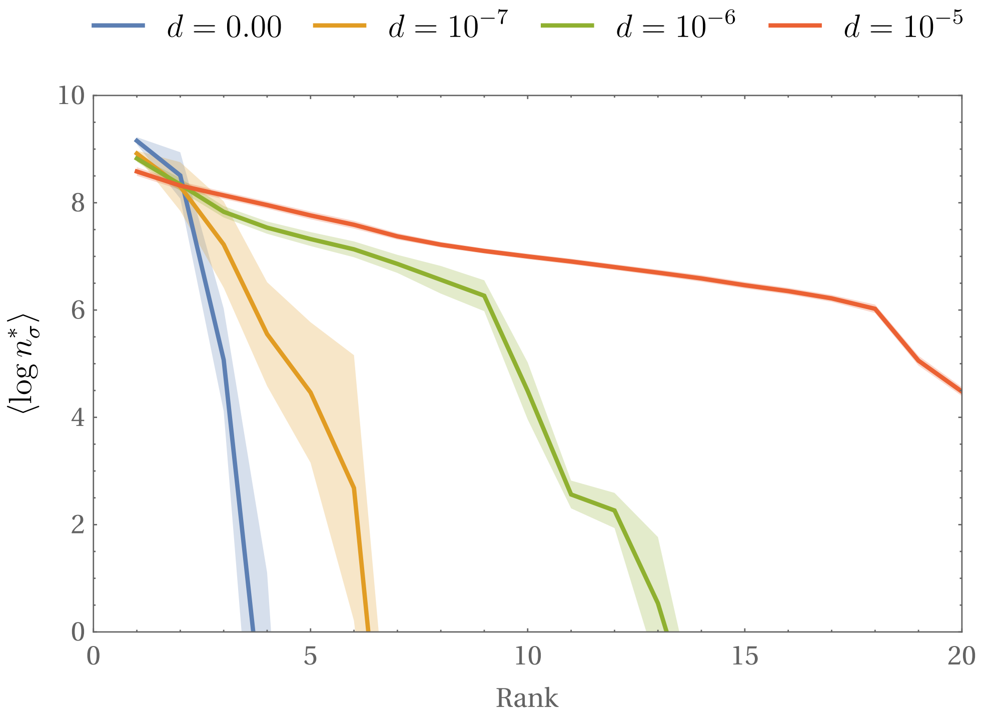


[p]

**FIG. S.9:** Rank distribution of the (decimal) logarithm of the stationary population densities for different adaptation velocities. We have used $m = 20$, $p = 3$, $\mathcal{Q} \in \mathcal{U}[10^{-7}, 5{\cdot}10^{-7}]$ g of resource/cells, $\delta_\sigma \in \mathcal{U}[5{\cdot}10^{-3}, 5{\cdot}10^{-2}]$ 1/h, $E_\sigma(0) \in \mathcal{U}[0, \mathcal{Q}\delta_\sigma]$, $v_i \in \mathcal{U}[0, 1]$ cells/g of resource. For each value of $d$ we have performed 100 iterations, each with its own $n_\sigma(0) \in \mathcal{U}[10^6, 5 \cdot 10^6]$ cells/mL, $c_i(0) \in \mathcal{U}[10^{-3}, 10^{-2}]$ g of resource/mL, $K_i \in \mathcal{U}[10^{-4}, 10^{-3}]$ g of resource/mL and the components of $\vec{s}$ drawn from $\mathcal{U}[10^{-3}, 10^{-2}]$ g of resource/(mL·h) so that it falls inside the convex hull of the initial metabolic strategies. Then, we have computed the logarithm of the stationary values of the species' populations (solving the equations until $T = 10^4$ h) and ordered them by rank; the opaque bands represent the standard error of the mean.

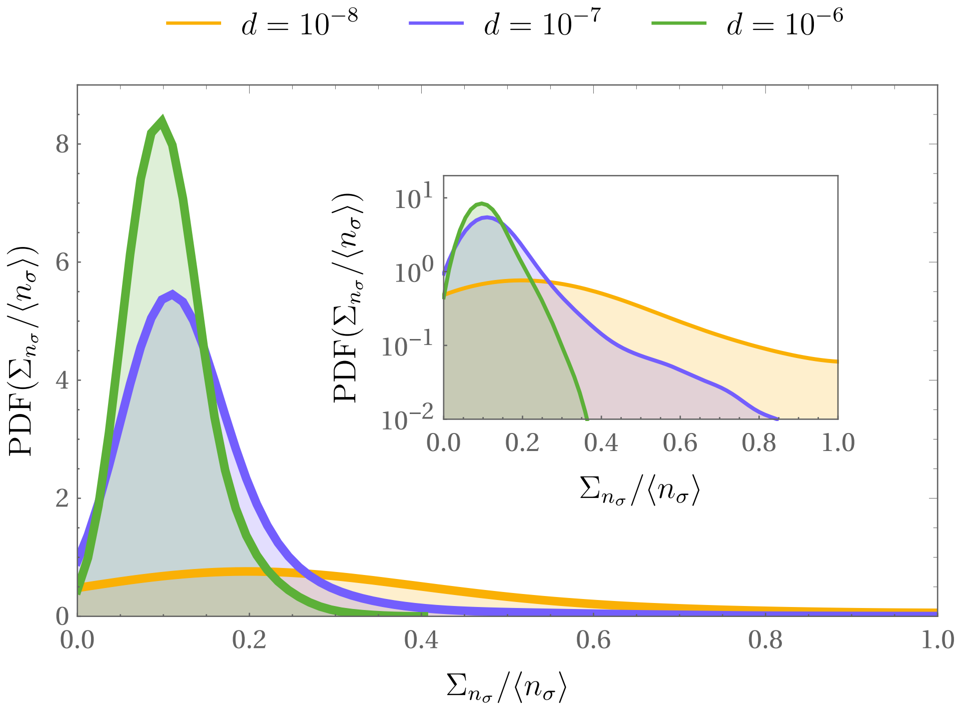


**FIG. S.10:** Probability density functions of the species' coefficients of variation $\Sigma_{n_\sigma}/\langle n_\sigma \rangle$ for different values of the adaptation velocity $d$. We have used $m = 20$, $p = 3$, $\mathcal{Q} \in \mathcal{U}[10^{-7}, 5 \cdot 10^{-6}]$ g of resoruce/cells, $\delta_\sigma \in \mathcal{U}[5 \cdot 10^{-3}, 5 \cdot 10^{-2}]$ 1/h, $E_\sigma(0) = \mathcal{Q}\delta_\sigma$, $v_i \in \mathcal{U}[10^8, 5 \cdot 10^9]$ cells/g of resource; $\vec{s}(t)$ is given as in (S.38) with $\tau_{\text{in}} = \tau_{\text{out}} = 48$ h. For each value of $d$ we have performed 1000 iterations, each with its own $n_\sigma(0) \in \mathcal{U}[10^6, 5 \cdot 10^6]$ cells/mL, $c_i(0) \in \mathcal{U}[10^{-3}, 10^{-2}]$ g of resource/mL, $K_i \in \mathcal{U}[10^-4, 10^-3]$ g of resource/mL and $\vec{s}_{\text{in}}$ and $\vec{s}_{\text{out}}$ chosen respectively inside and outside the convex hull of the initial metabolic strategies, and we have computed $\Sigma_{n_\sigma}/\langle n_\sigma \rangle$ as described in III E with $T = 10^4$ h. The PDFs have then been computed using kernel density estimation (we have used Mathematica's `SmoothKernelDistribution` function). As we can see, as $d$ grows the species' coefficients of variation are increasingly concentrated around smaller and smaller values, i.e. the species' populations become more stable as adaptation accelerates. **Inset**: Same plot with logarithmic scale on the vertical axis.
