## Supplementary figures and images for "Adaptive metabolic strategies in consumer-resource models"

### Animation of figure S4

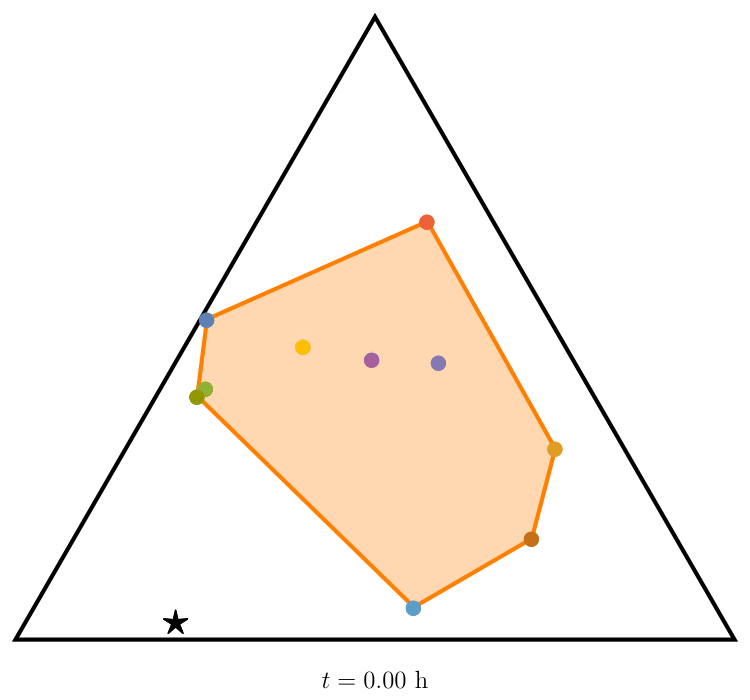

### Animation of figure S5

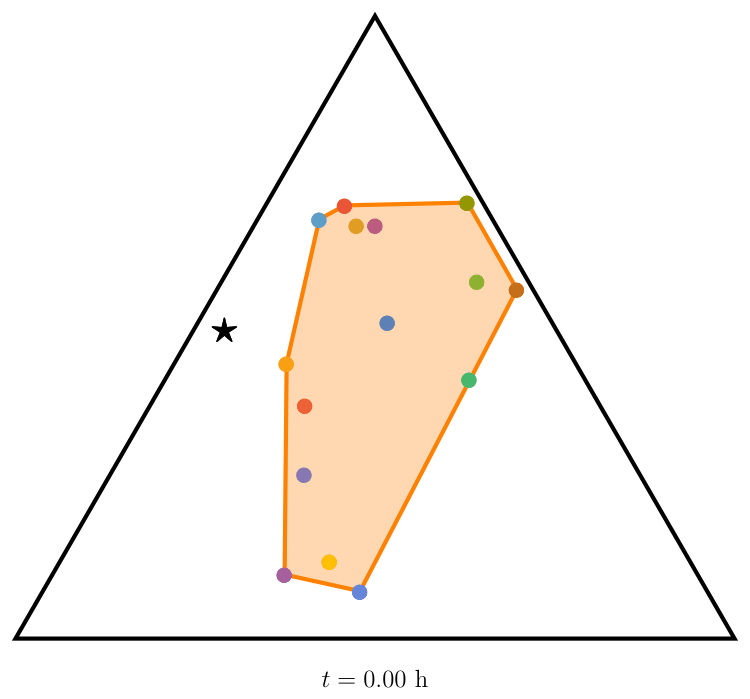

### Animation of figure S6a and S6b

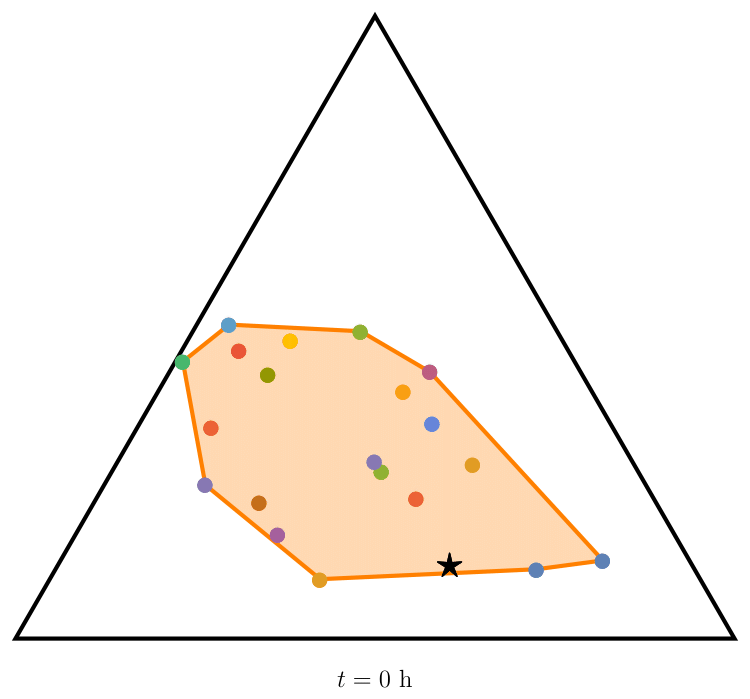

### Animation of figure S6a and S6d

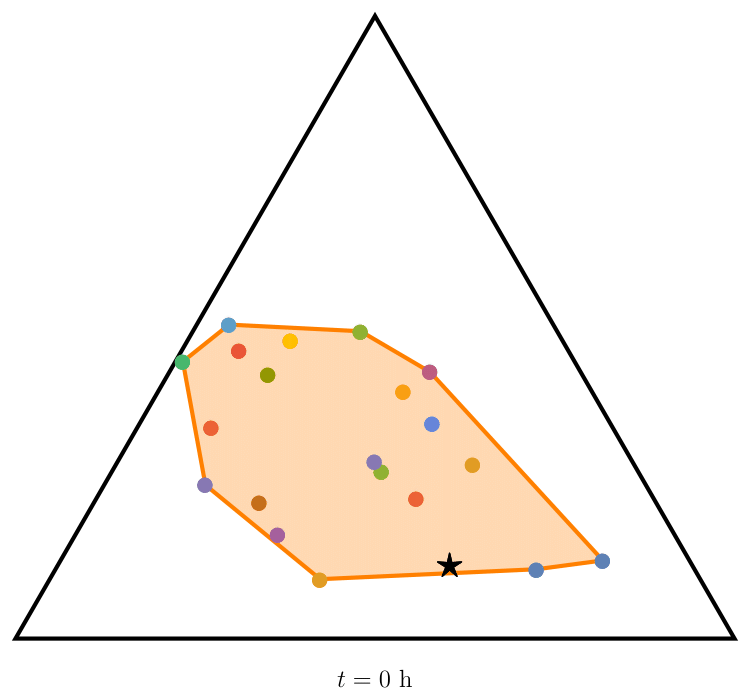

### Animation of figures 3 and S3

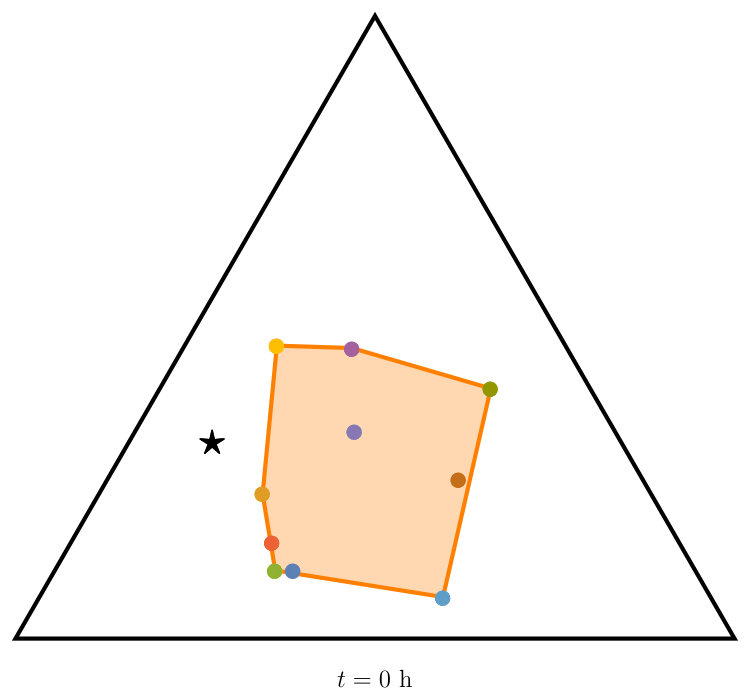
